# A metabolically stable PET tracer for imaging synaptic vesicle protein 2A: Synthesis and preclinical characterization of [^18^F]SDM-16

**DOI:** 10.1101/2021.06.25.449978

**Authors:** Chao Zheng, Daniel Holden, Ming-Qiang Zheng, Richard Pracitto, Kyle C. Wilcox, Marcel Lindemann, Zachary Felchner, Li Zhang, Jie Tong, Krista Fowles, Sjoerd J. Finnema, Nabeel Nabulsi, Richard E. Carson, Yiyun Huang, Zhengxin Cai

## Abstract

**Purpose:** To investigate the synaptic vesicle glycoprotein 2A (SV2A) expression in the whole central nervous system and peripheral tissues, a metabolically stable SV2A radiotracer is desirable to minimize a potential confounding effect of radiometabolites. The aim of this study was to develop and evaluate a metabolically stable SV2A radiotracer, [^18^F]SDM-16, in nonhuman primate brains.

**Methods:** The racemic SDM-16 (4-(3,5-difluorophenyl)-1-((2-methyl-1H-imidazol-1-yl)methyl)pyrrolidin-2-one) was synthesized and assayed for *in vitro* SV2A binding affinity. We synthesized the enantiopure [^18^F]SDM-16 using the corresponding arylstannane precursor. Nonhuman primate brain PET was performed on a FOCUS 220 system. Arterial blood was drawn for metabolite analysis and construction of plasma input function. Regional time-activity curves (TACs) were evaluated with the one-tissue compartment (1TC) model to obtain the volume of distribution (*V*_T_). Binding potential (*BP*_ND_) was calculated using either the nondisplaceable volume of distribution (*V*_ND_) or the centrum semiovale (CS) as the reference region.

**Results:** Racemic SDM-16 was synthesized in 3 steps with 44% overall yield and has high affinity (*K*_i_ = 3.7 nM) to human SV2A. [^18^F]SDM-16 was prepared in greater than 99% radiochemical and enantiomeric purity. This radiotracer displayed high specific binding in brain and was metabolically more stable than other SV2A PET tracers. The plasma free fraction (*f*_P_) of [^18^F]SDM-16 was 69%, which was higher than those of [^11^C]UCB-J (46%), [^18^F]SynVesT-1 (43%), [^18^F]SynVesT-2 (41%), and [^18^F]UCB-H (43%). The TACs were well described with the 1TC. The averaged test-retest variability (TRV) was −9±8%, and averaged absolute TRV (aTRV) was 10±7% for all analyzed brain regions.

**Conclusion:** We have successfully synthesized a metabolically stable and high affinity SV2A PET tracer, [^18^F]SDM-16, which showed high specific and reversible binding in the NHP brain. [^18^F]SDM-16 may have potential application in the visualization and quantification of SV2A beyond the brain.

## INTRODUCTION

Proteins in the synaptic vesicle glycoprotein 2 (SV2) family located in presynaptic terminals are essential components of synaptic vesicles [1]. As one of the isoforms, SV2A is ubiquitously expressed in virtually all synapse terminals, and involved in the regulation of synaptic exocytosis and endocytosis [2, 3]. SV2A is a known target for anti-epilepsy drugs, such as levetiracetam (Keppra^®^, LEV) [4]. Positron emission tomography (PET) is a non-invasive quantitative imaging modality that provides functional and physiological information in living systems. SV2A PET tracers can be used to study receptor occupancy in clinical development of new drug candidates targeting SV2A, and to measure changes of SV2A in neuropsychiatric diseases [5-10]. SV2A PET has potential applications beyond the brain, as SV2A is expressed not only in the central nervous system (CNS) [11], but also in neuroendocrine cells, ganglia cells in the peripheral nervous system (PNS) [12, 13], and enriched in several types of cancers [14]. While the current metabolically labile SV2A PET tracers are suitable for brain PET imaging due to the blood-brain barrier (BBB) preventing their radiometabolites from entering the brain, a more metabolically stable radiotracer is desirable for the investigation of SV2A expression in other organs and relationship between SV2A expression in CNS and PNS, to minimize the confounding effect of radiometabolites.

Several SV2A PET tracers have been synthesized and evaluated in animals and human during the past few years by our group and others (**Fig. 1**) [7, 15]. [^18^F]UCB-H (**2**) [16-18] was the first SV2A PET tracer tested in human [17], followed by [^11^C]UCB-J (**3**) [5], [^11^C]UCB-A (**1**), [^18^F]SynVesT-1(a.k.a. [^18^F]SDM-8 or [^18^F]MNI-1126) (**6**) [19, 20], and [^18^F]SynVesT-2 (a.k.a. [^18^F]SDM-2) (**7**) [21]. The isotopologue of **3**, [^18^F]UCB-J was evaluated in rhesus monkeys, but not pursued for clinical evaluation [22]. [^11^C]UCB-J is currently the SV2A PET tracer most widely used in PET imaging investigations of neuropsychiatric disorders, i.e., epilepsy, Alzheimer’s disease, Parkinson’s disease, schizophrenia, major depressive disorder, and posttraumatic stress disorder [5, 23-25]. Among the available SV2A PET tracers, [^11^C]UCB-A was arguably the most metabolically stable, even though its prevalent radiometabolite species in the plasma were not identified yet [26, 27]. However, the relatively short radioactive half-life (~ 20 min) of carbon-11 together with the relatively slow kinetics in the brain limited the potential clinical application of [^11^C]UCB-A [28, 29]. We hypothesized that the slow kinetics of [^11^C]UCB-A (as reflected in its low *K*_1_ and *k*_2_ values) was due to its relatively low membrane permeability, which is associated with its low hydrophobicity (LogD: 1.1). To develop a metabolically stable analog of [^11^C]UCB-A combined with an improved pharmacokinetic (PK) profile, we synthesized and evaluated a novel ^18^F-labeled SV2A PET tracer, [^18^F]SDM-16 (**7**), in nonhuman primates.

**Figure 1:**
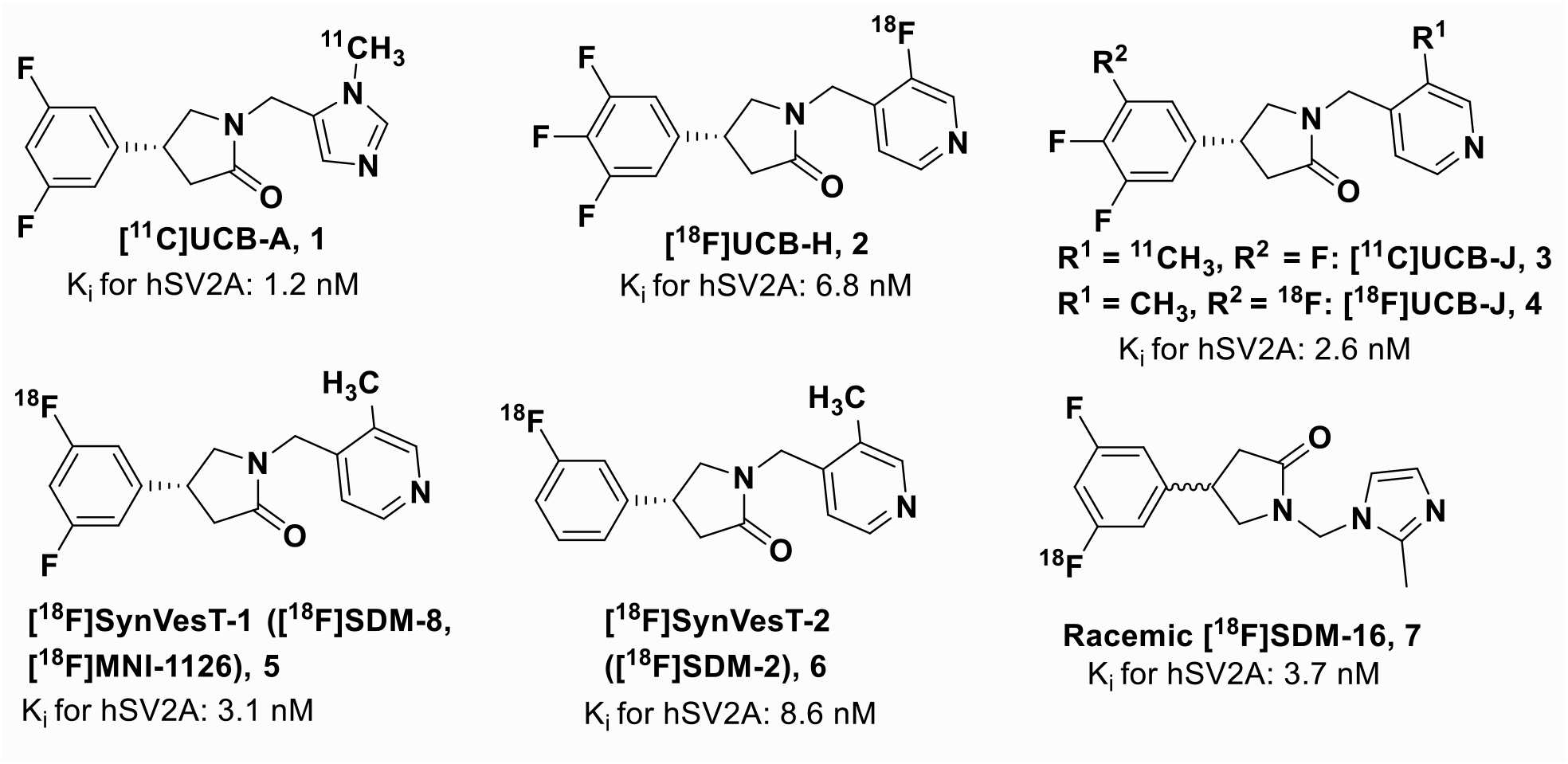
Current SV2A radiotracer

## MATERIALS AND METHODS

### Chemistry

All compounds were prepared from commercially available starting materials. Details are described in the supplemental materials.

### Preparation of (*R*)-4-(3-fluoro-5-(fluoro-^18^F)phenyl)-1-((2-methyl-1*H*-imidazol-1-yl)methyl)pyrrolidin-2-one ([^18^F]SDM-16)

The cyclotron produced aqueous [^18^F]fluoride solution in H_2_^18^O was transferred to a V-vial in a lead-shielded hot cell, where the [^18^F]fluoride anion was trapped on an anionic exchange resin cartridge (Chromafix-PS-HCO_3_) pre-activated by elution sequentially with EtOH (5 mL), an aqueous solution of potassium triflate (KOTf, 90 mg/mL, 5 mL), and deionized (DI) water (5 mL). The potassium [^18^F]fluoride was then eluted off the cartridge into a 2 mL V-vial with the mixture of aqueous solutions of KOTf (10 mg/mL, 0.45 mL) and K_2_CO_3_ (1 mg/mL, 50 μL), and MeCN (0.5 mL). The eluent was azeotropically dried at 110 °C, with two portions of anhydrous MeCN (1.0 mL × 2) added during the process. A solution of the precursor **17** (1.7-3.0 mg) in anhydrous *N*,*N*-dimethylacetamide (DMA, 0.4 mL) was then added to the reaction vial, followed by the solution of pyridine (1 M in DMA, 0.1 mL) and copper(II) triflate (0.2 M in DMA, 67 μL). The reaction mixture was then heated at 110 °C for 20 min, diluted with the HPLC mobile phase (1.5 mL) and purified by HPLC (column: Genesis C18, 4 μm, 10 × 250 mm; mobile phase: 19% CH_3_CN and 81% 0.1 M ammonium formate solution with 0.5% AcOH, pH 4.2; flow rate: 5 mL/min). The eluent was monitored by a UV detector (at 254 nm) and a radioactivity detector. The fraction containing [^18^F]SDM-16 was collected, diluted with DI water (50 mL), and passed through a C18 SepPak (No. 50-819-184, Waters), which was then washed with 0.001N HCl (10 mL) and dried with 10 ml air. The product was eluted off with EtOH (1 mL), diluted with USP grade saline (3 mL), passed through a sterile membrane filter (0.22 μm), and collected in a sterile vial pre-charged with 7 mL of USP saline and 20 μL of 8.4% NaHCO_3_ to afford a formulated solution ready for administration.

### Competition Radioligand Binding Assay

Competition binding assays were performed twice in independent experiments. The racemic SDM-16 standard compound was dissolved in DMSO (10 mM), which was diluted in PBS pH 7.4 (Gibco) with 0.1% BSA assay buffer to give 12 half-log dilutions from 10 μM to 32 pM. Duplicate samples of human frontal cortex gray matter were homogenized in PBS buffer (10 mg/mL) for storage at −80 °C and were diluted to a stock concentration of 4 mg/mL in PBS on the day of the assays. [^3^H]UCB-J was obtained with a molar activity of 1.29 TBq/mmol (34.9 Ci/mmol) and radiochemical purity of 98.9%, diluted in duplicate to a stock concentration of 6.25 nM. Working stocks of brain homogenate (100 μL; final concentration of 2 mg/mL), blocking ligands (20 μL), and radioligand (80 μL; final concentration 2.5 nM) were combined in quadruplicate wells of 96-well plates, sealed, and incubated at room temperature for 90 minutes on an orbital shaker set to 250 RPM. Reaction plates were filtered, rapidly washed with cold PBS, and dried. 40 μL Microscint-20 scintillation cocktail (Perkin-Elmer) was added to each well and the plate was counted using a Microbeta2 plate reader (Perkin-Elmer). GraphPad Prism was used for curve fitting using the one-site *K*_i_ model.

### Measurement of Lipophilicity

The logP of [^18^F]SDM-16 was determined by a method modified from previously published procedures [30]. Briefly, an aliquot of 70 kBq (10 μCi) of the radioligand was added to a 2 mL microtube containing 0.8 mL of octanol and 0.8 mL of 1× phosphate buffered saline (1× PBS, pH 7.4). The mixture was vortexed for 30 s and then centrifuged at 2000 g for 2 min. A subsample of the octanol (0.1 mL) and 1× PBS (0.5 mL) layers was evaluated with a gamma counter. The major portion of the octanol layer (0.5 mL) was diluted with another 0.3 mL of octanol, mixed with a fresh portion of 0.8 mL of PBS, vortexed, centrifuged, and analyzed as described above. This process was repeated until consistent log *P* values were obtained, with five consecutive equilibration procedures being performed for each logP measurement. Four separate measurements were performed for [^18^F]SDM-16 on different days.

### Measurement of Plasma Free Fraction (*f*_p_)

The unbound fraction of [^18^F]SDM-16 in plasma (*f*_p_) of rhesus monkey was measured in triplicate using the ultrafiltration method [19, 21]. Briefly, [^18^F]SDM-16 solution was added to 3 mL of whole blood. After incubation at ambient temperature for 5 min, the blood sample was centrifuged at 3900 rpm for 5 min. A sample of the supernatant plasma (0.3 mL) was loaded onto the reservoir of a Centrifree^®^ Ultrafiltration device (Merck Millipore Ltd. Tullagreen, Carrigtwohill, Co. Cork, IRELAND) in triplicate and centrifuged at 1228 g for 20 min. The *f*_p_ value was calculated as the ratio of radioactivity in the filtrate to that in the plasma.

### PET Imaging Experiments in Rhesus Monkeys

A total of 5 PET imaging experiments with [^18^F]SDM-16 were performed in rhesus monkeys (*Macaca mulatta*) according to a protocol approved by the Yale University Institutional Animal Care and Use Committee (IACUC). Two monkeys were studied. One monkey (8 years old, Male, 9.5 kg) underwent two baseline scans and one blocking scan and the other monkey (13 years old, Female, 9.5kg) underwent one displacement scan and one whole-body scan. Rhesus monkeys were fasted overnight and sedated using intramuscular injection of alfaxalone (2 mg/kg), midazolam (0.3 mg/kg), dexmedetomidine (0.01 mg/kg), and anaesthetized with 0.75-2.5% isoflurane approximately 2 h before the PET scan. Anesthesia was subsequently maintained with isoflurane (1.5%-2.5%) for the duration of the imaging experiments. Body temperature was maintained by a water-jacket heating pad. The animal was attached to a physiological monitor, and vital signs (heart rate, blood pressure, respirations, SPO_2_, EKG, ETCO_2_, and body temperature) were continuously monitored. A venous line was inserted in one limb for administration of radiotracer, displacement and blocking drugs. A catheter was placed in the femoral artery in the other limb for blood sampling. Dynamic PET brain scans were performed on a Focus 220 system (Siemens Medical Solutions, Knoxville, TN, USA) with a reconstructed image resolution of approximately 1.5 mm. After a 9 min transmission scan, the radioligand was injected i.v. over 3 min. by an infusion pump. Dynamic PET scans were performed for three hours (baseline and blocking scans) or four hours (displacement scan). For the blocking scan LEV (30 mg/kg) was administered intravenously at 10 min before tracer injection, while in the displacement scan the same dose of LEV was infused at 120 min after tracer injection.

PET images were reconstructed with built-in corrections for attenuation, normalization, scatter, randoms, and deadtime. PET brain images were registered to the animal’s MR image, which was subsequently registered to a brain atlas to define the regions of interest. Dynamic images were reconstructed using a Fourier rebinning and filtered back projection algorithm. A rhesus monkey brain atlas was used for generation of regions of interest (ROIs) and time–activity curves (TACs) for the following ROIs: amygdala, brain stem, caudate nucleus, centrum semiovale (CS), cerebellum, cingulate cortex, frontal cortex, globus pallidus, hippocampus, insula, nucleus accumbens, occipital cortex, pons, putamen, substantia nigra, temporal cortex, and thalamus.

### Plasma Radiometabolite Analysis

Arterial blood samples were collected during the PET scans to measure the radioactivity in plasma for generation of the metabolite-corrected arterial plasma input function. Plasma radiometabolite analysis was performed using the column switching method, following a published protocol [31]. Briefly, arterial blood samples were collected at 3, 8, 15, 30, 60, 90, 120 and 180 min post-injection (p. i.), treated with urea (8 M), filtered, and injected onto a self-packed short column (4.6 × 19 mm) eluting with 1% MeCN in water at a flow rate of 2 mL/min. The sample was then back flushed onto a Gemini-NX column (5 μm, 4.6 mm × 250 mm) eluting with 40% MeCN/60% 0.1 M ammonium formate (pH 6.4) at a flow rate of 1.2 mL/min. The eluent was fraction-collected using an automated Spectrum Chromatography CF-1 fraction collector. Activity in the whole blood, plasma, filtrated plasma-urea mix, filters, and HPLC fractions were counted with automatic gamma well-counter (Wizard 2, PerkinElmer). The sample recovery rate, extraction efficiency, and HPLC fraction recovery were monitored. The unmetabolized [^18^F]SDM-16 parent fraction was determined as the ratio of the sum of radioactivity in fractions containing the parent compound to the total amount of radioactivity collected and fitted with inverted Gamma4 approaches.

### Kinetic Modeling

Volume of distribution (*V*_T_, mL·cm^−3^) values and the first-order kinetic rate constants of tracer (*K*_1_) were derived through 1-tissue (1T) compartment kinetic modeling with the metabolite-corrected arterial plasma input function, which was calculated as the product of the fitted total plasma curve and the parent radiotracer fraction curve. Nondisplaceable volume of distribution (*V*_ND_) and SV2A occupancy by LEV was calculated using the Lassen plot [32]. Nondisplaceable binding potential (*BP*_ND_) values were calculated from *V*_T_ values using CS as reference region, or the *V*_ND_ derived from the blocking study, i.e., *BP*_ND_ = (*V*_T, ROI_ – *V*_T, CS_)/*V*_T, CS_ or *BP*_ND_ = (*V*_T_/*V*_ND_) −1.

### Radiation Dosimetry Study

A whole-body biodistribution study was performed in one rhesus monkey (Female, 9.4 kg) to estimate human organ radiation dosimetry. The scan was carried out on a Biograph mCT (hybrid PET/CT, Siemens Medical Systems, Knoxville, TN) scanner following an i.v. injection of 187.6 MBq (5.1 mCi) [^18^F]SDM-16 and a mass dose of 0.27 μg at time of injection. The monkey was scanned for about 4 h in a sequence of 22-24 passes from top of the head to the mid-thigh. Scans were reconstructed and visually inspected for organ activity concentrations exceeding background level. The organs included were brain, heart, liver, gall bladder, spleen, kidneys, and urinary bladder contents. Regions of interest were delineated on these organs and mean activity values were computed to form TACs.

Within-pass decay correction was removed to reflect the actual activity in each organ, and cumulative activity (Bq·h/cm^3^) computed by integration of the data from the scan. The tail portions beyond the end of the scan were extrapolated assuming only physical decay of the tracer. These values were multiplied by the organ volumes of a standard 55 kg adult female reference mathematical phantom, and then normalized to injected activity to obtain organ residence times (N, h). Final values were then entered into the OLINDA software to obtain absorbed doses in all organs, which were computed with different voiding assumptions.

## RESULTS

### Chemistry

Compounds **9** and **15** were synthesized from commercially available aldehydes **8** and **14**, respectively in 3 steps as racemic mixtures (**Schemes 1** and **2**), following the published procedures [19], with minor modifications. After chiral resolution of the racemic (***rac***) products, (*R*)-**9** and (*R*)-**15** were obtained with enantiomeric excess (*e.e*.) greater than 99%. Their absolute configuration was determined by X-ray crystallography. Condensation of compound **9** with formalin, followed by substitution of the intermediate **10** with 2-methyl-1*H*-imidazole afforded ***rac*** SDM-16 in 44% overall yield. Condensation of 2-methyl-1*H*-imidazole with paraformaldehyde gave (2-methyl-1*H*-imidazol-1-yl)methanol (**11**), which was chlorinated by thionyl chloride to give the imidazole salt **12**. Nucleophilic substitution of chloride **12** with (*R*)-**19** or (*R*)-**15**, gave SDM-16 standard **13** or the bromo analog **16** in 92% and 90% yield, respectively. It is worth mentioning that when the chlorine atom in **12** was replaced with other leaving groups (Br, OTs, OTf), the substitution reactions did not yield the desired product. As the radiolabeling precursor, arylstannane **17**, was obtained from **16** in 47% yield via Pd(0)-catalyzed stannylation reaction. Finally, [^18^F]SDM-16 (**18**) was prepared from the enantiopure precursor **17** with > 99.9% radiochemical and enantiomeric purity, as determined by reverse phase C18 and chiral HPLC analysis. Molar activity at the end of synthesis (EOS) was 283 ± 42 GBq/μmol (n = 6).

**Scheme 1:**
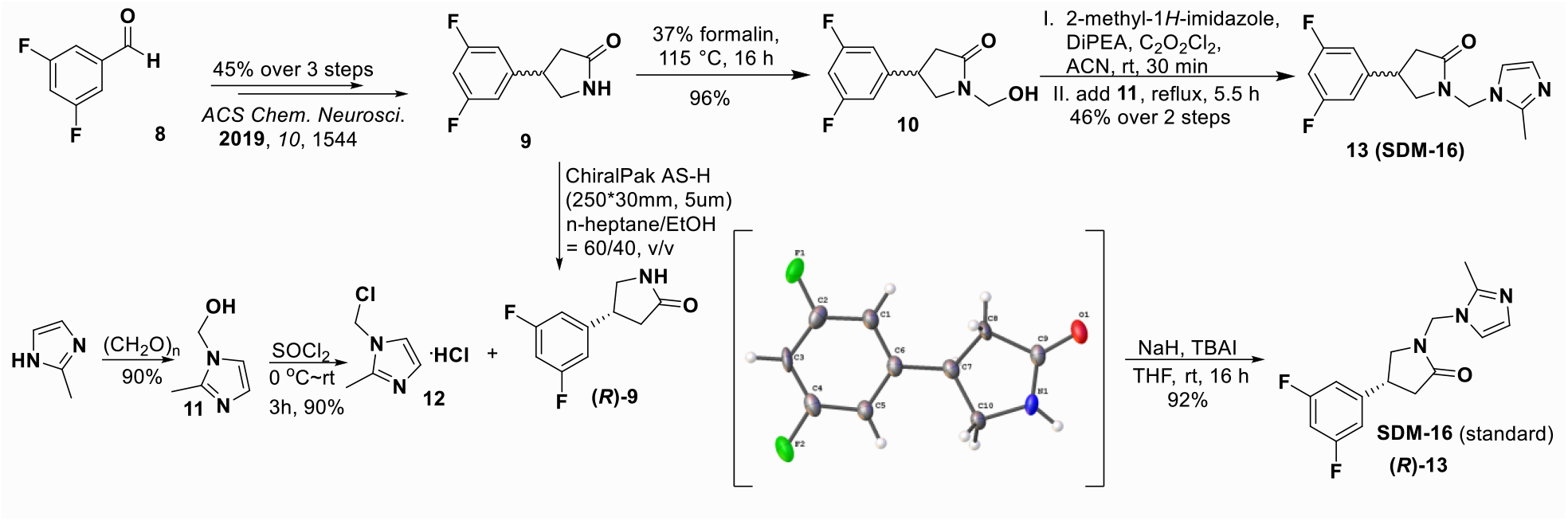
Synthesis of the racemic and enantiopure standard compound SDM-16 (**13**).

**Scheme 2:**
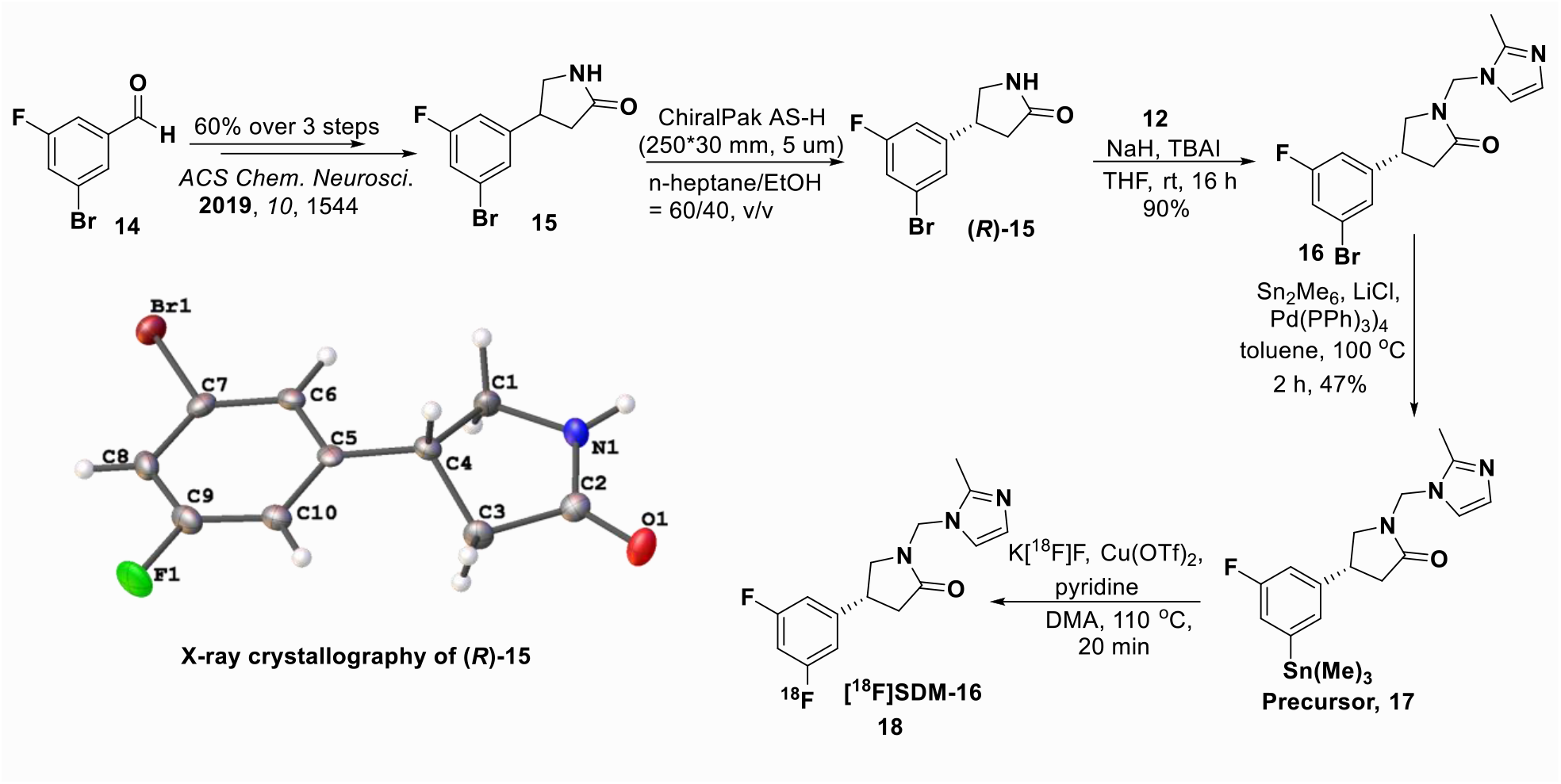
Synthesis of the [^18^F]SDM-16 (18) and its enantiopure labeling precursor (**17**).

### In Vitro Competition Binding Assay

Racemic SDM-16 possessed high binding affinity to human SV2A, with *K*_i_ of 3.7 nM in our radioligand competition binding assay using [^3^H]UCB-J and human frontal cortex tissue homogenate. For comparison, *K*_i_ values were 2.6 nM, 3.1 nM and 8.6 nM for UCB-J, SynVesT-1 and SynVesT-2 (**Fig. 1**) in the same assay [33].

### Measurement of Lipophilicity

The averaged LogP value of [^18^F]SDM-16 was 1.65 ± 0.05 (n = 20), which was lower than that of [^11^C]UCB-J (2.46), [^18^F]UCB-H (2.31), and [^18^F]SynVesT-1 (2.32), and higher than [^11^C]UCB-A (1.10), and within the optimal range for BBB penetration (1 < LogP < 3) [34].

### PET Imaging Experiments in Rhesus Monkeys

The injected radioactivity ranged from 183 to 188 MBq (n = 5), corresponding to 0.646 – 0.926 μg of SDM-16. At this microdose level, no adverse events were observed throughout the imaging study. Adverse event was also not observed following LEV (i.v., 30 mg/kg) administration, including the displacement and blocking scan.

#### Plasma Analysis

After the administration of [^18^F]SDM-16, the tracer concentration in the plasma showed a sharp increase within 5 min, followed by a fast distribution phase and a slow clearance phase. [^18^F]SDM-16 had higher metabolite-corrected plasma SUV than the other SV2A radiotracers, indicating slower plasma clearance and higher metabolic stability (**Fig. 2a**). Whole blood and plasma-input functions were highly consistent between the two animals, with a stable plasma to whole-blood ratio of 0.93 ± 0.13 over the entire 180-min acquisition period (**Fig. S1**). In rhesus monkeys, [^18^F]SDM-16 was metabolized slowly, with 88 ± 8% and 81 ± 8% intact radiotracer present in the plasma at 30 and 120 min post-injection (p.i., n = 4, **Fig. 2b**), respectively, compared with 70 ± 7%, 42 ± 13%, 40 ± 6%, and 30 ± 3% parent fraction at 30 min p.i. for [^11^C]UCB-A (n = 5), [^18^F]SynVesT-1 (n = 5), [^11^C]UCB-J (n = 11), and [^18^F]UCB-H [35], respectively. All observed radiometabolite fractions in the plasma had shorter retention times than the parent tracer, indicating that they were more hydrophilic and less likely to penetrate the BBB (**Fig. 2c**).

**Figure 2:**
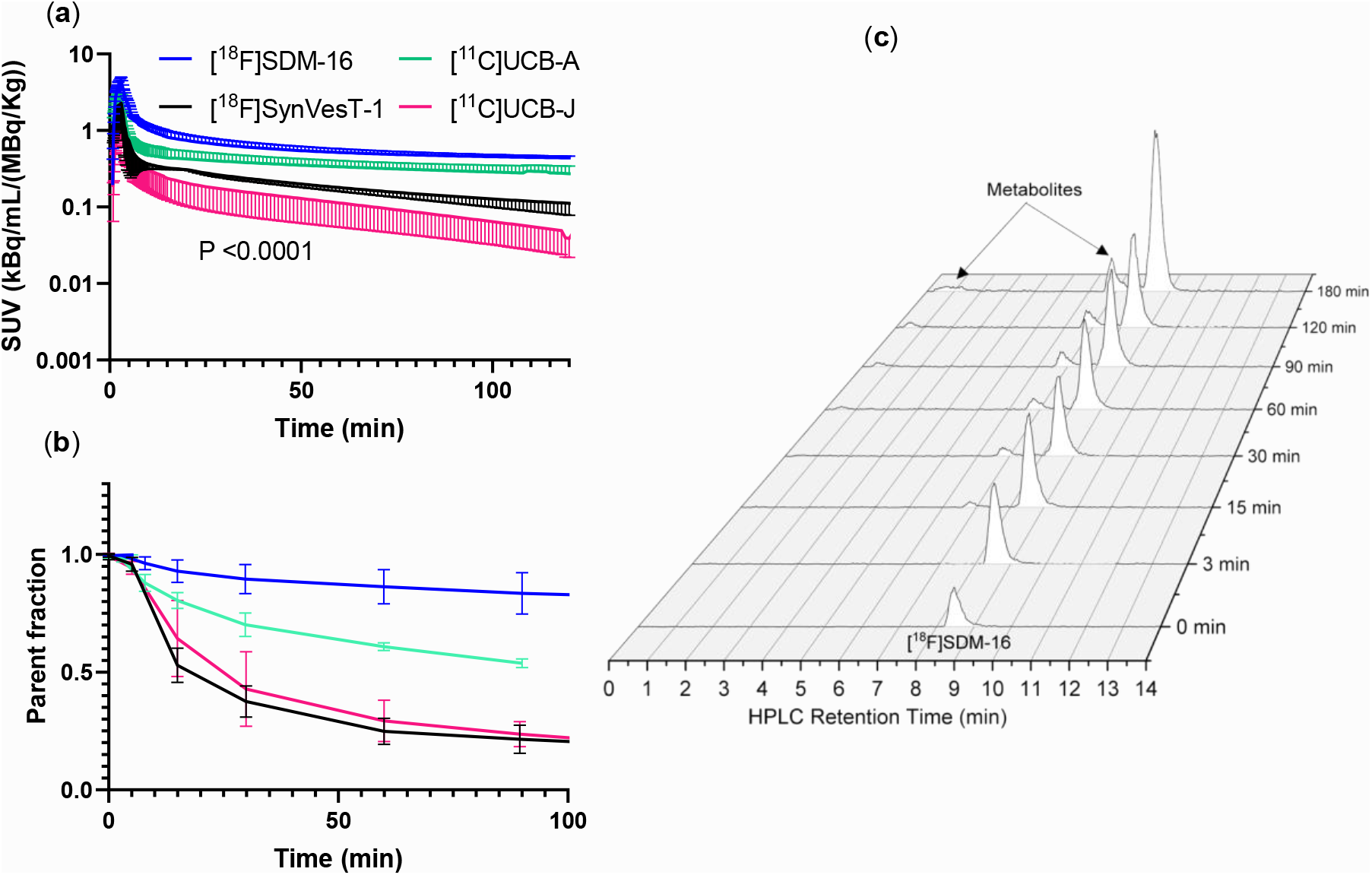
Plasma analysis of the 4 radiotracers in monkey. (a) Metabolite-corrected plasma activity for [^18^F]SDM-16 (n = 4), [^18^F]SynVesT-1 (n = 2), [^11^C]UCB-A (n = 6), and [^11^C]UCB-J (n = 5), with 2-way ANOVA analysis; (b) plasma parent fraction over time for [^18^F]SDM-16 (n = 4), [^18^F]SynVesT-1 (n = 5), [^11^C]UCB-A (n = 4), and [^11^C]UCB-J (n = 5); (c) radio-HPLC chromatograms of plasma metabolite analysis of [^18^F]SDM-16 (retention time at 8.5 min). The retention time of the major radiometabolite was 6.8 min and a minor radiometabolite at around 0.5-1.2 min.

#### Brain PET image analysis

Summed SUV images from the baseline and blocking scans of [^18^F]SDM-16 are shown in **Fig. 3**. At baseline, high contrast between gray matter and ventricles was clearly visible (**Fig. 3a**); while blocking with LEV significantly reduced the tracer uptake in grey matters (**Fig. 3b**). [^18^F]SDM-16 had an apparently slow kinetic profile, with tracer uptake increasing gradually till the end of the scan to an SUV of about 10 (for frontal cortex and putamen, **Fig. 3c**), which was higher than for [^11^C]UCB-A (SUV about 4 in **Fig. 3d**). Nevertheless, the binding of [^18^F]SDM-16 was reversible, as demonstrated by the LEV displacement experiment, in which the tracer uptake was reduced by 40.5 ± 0.1% (averaged from 5 brain regions), based on the SUV values at the end of the displacement scan (4 h p.i.) normalized by those before the administration of LEV (**Fig. 3e**). In the blocking study, the preinjected LEV (i.v., 30 mg/kg), resulted in 57±10% reduced tracer uptake in gray matter regions, based on the normalized terminal SUV values in the blocking scan to those of the baseline scans (**Fig. 3f**). Our displacement and pre-blocking PET imaging results confirmed the reversible and SV2A-specific binding of [^18^F]SDM-16 in nonhuman primates. We did not observe any radioactivity in skull throughout the PET imaging window (up to 4 h p.i.), indicating the lack of *in vivo* defluorination.

**Figure 3:**
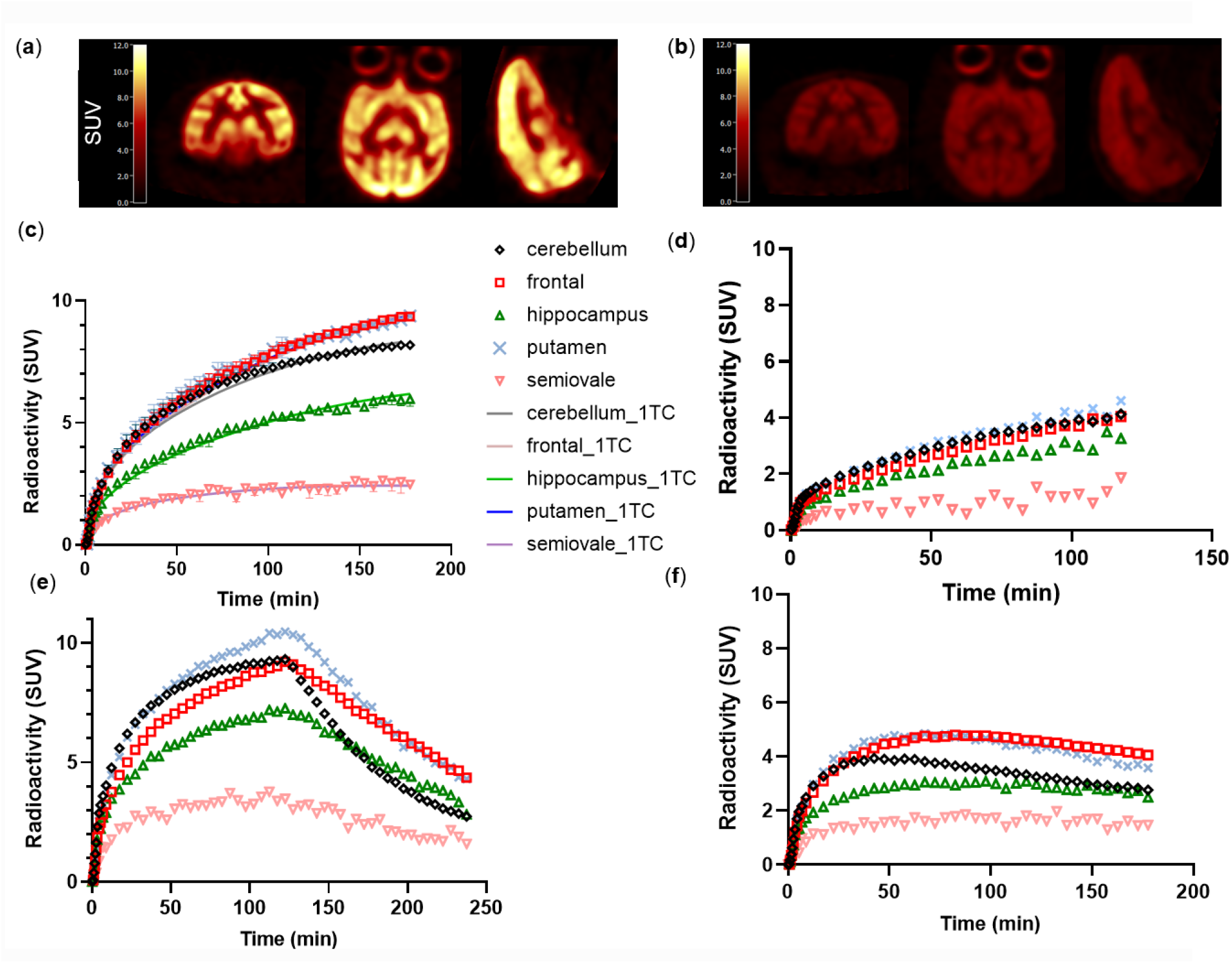
Summed SUV images of [^18^F]SDM-16 in the brain of a rhesus monkey from 150 to180 min imaging window at (a) baseline scan and (b) blocking scan with LEV (30 mg/kg, i.v.). Time activity curves of (c) baseline scans of [^18^F]SDM-16, (d) baseline scan of [^11^C]UCB-A, (e) displacement study of [^18^F]SDM-16, with LEV given at 120 min p.i., and (f) pre-blocking study of [^18^F]SDM-16 with LEV (30 mg/kg, i.v.).

#### Kinetic modeling

Regional time-activity curves (TACs) were fitted with 1-tissue compartment (1TC) model to generate binding parameters, using the metabolite-corrected plasma input function. Similar to [^11^C]UCB-J, [^18^F]SynVesT-1 and [^18^F]SynVesT-2, the 1TC model described the TACs well (**Fig. 3c**) and provided reliable estimates of regional volumes of distribution (*V*_T_) for [^18^F]SDM-16. The [^18^F]SDM-16 *V*_T_ values were highest in cingulate cortex (40.6 mL/cm^3^), followed by caudate, putamen, and thalamus, and lowest in centrum semiovale (CS, 6.4 mL/cm^3^) (**Table 1**). The rank order of 1TC model generated *V*_T_ values was consistent with previously reported SV2A PET tracers and *in vitro* binding results [4, 19], and the monkey brain *V*_T_ values of [^18^F]SDM-16 correlated well with those of [^11^C]UCB-A (Y = 1.19*X + 5.10, R^2^ = 0.80, p = 0.0001), [^18^F]SynVesT-1 (Y = 1.06*X + 5.62, R^2^ = 0.88, p = 0.0001) and [^11^C]UCB-J (Y = 1.23*X + 8.27, R^2^ = 0.95, p = 0.0001) (**Fig. 4**) [36]. Because different monkeys were used in the evaluations of these SV2A PET tracers, we observed more variability in these plots than what has previously been shown in plots using data from the same subject [36].

**Table 1:** 1TC-derived regional *V*_T_ values (mean ± SD) from baseline scans (n=2) and blocking (n=1) of [^18^F]SDM-16, along with those from baseline scans of [^18^F]SynVesT-1 (n=3) [19], [^11^C]UCB-J (n=5) [26], and [^11^C]UCB-A (n =5).

| | $V_T$ (baseline) | | | | $V_T$ (blocking) |
| --- | --- | --- | --- | --- | --- |
| | [$^{11}\text{C}$]UCB-A | [$^{11}\text{C}$]UCB-J | [$^{18}\text{F}$]SynVesT-1 | [$^{18}\text{F}$]SDM-16 | [$^{18}\text{F}$]SDM-16 |
|  | (n=5) | (n=5) | (n=3) | (n=2) | (n=1) |
| Cingulate cortex | 54.96±24.29 | 55.57±9.94 | 48.5±8.9 | 40.57±0.72 | 10.43 |
| Frontal cortex | 48.96±18.24 | 55.36±8.18 | 47.1±9.1 | 34.85±1.93 | 9.52 |
| Insular cortex | 57.78±26.87 | 54.56±6.68 | 46.1±9.2 | 37.52±4.90 | 10.27 |
| Nucleus accumbens | 34.13±4.75 | 53.94±9.18 | 45.2±7.9 | 39.39±0.58 | 10.44 |
| Occipital cortex | 47.04±16.55 | 52.90±7.03 | 43.7±10.7 | 34.60±2.76 | 8.32 |
| Temporal cortex | 51.41±26.22 | 50.45±6.89 | 42.2±10.0 | 33.17±1.38 | 8.62 |
| Putamen | 48.78±18.76 | 45.20±3.04 | 34.6±4.9 | 31.77±2.07 | 8.81 |
| Caudate | 36.61±13.90 | 44.81±4.59 | 35.0±6.3 | 24.79±1.73 | 7.68 |
| Thalamus | 31.92±10.11 | 40.26±6.44 | 34.2±3.0 | 26.58±0.47 | 8.24 |
| Cerebellum | 30.73±9.94 | 39.54±5.82 | 28.9±6.8 | 26.60±1.65 | 6.60 |
| Hippocampus | 31.44±16.65 | 34.43±2.78 | 30.1±5.6 | 19.17±2.47 | 5.89 |
| Globus pallidus | 32.18±12.17 | 27.94±2.96 | 21.6±4.2 | 19.30±0.83 | 6.93 |
| Brainstem | 21.03±4.94 | 23.52±2.84 | 16.3±3.6 | 15.06±0.38 | 4.54 |
| Amygdala | 15.62±8.2 | 24.43±1.98 | 24.8±8.5 | 9.97±0.64 | 4.46 |
| Centrum semiovale | 10.88±5.14 | 13.59±2.85 | 8.9±2.0 | 6.40±0.37 | 3.35 |

**Figure 4:**
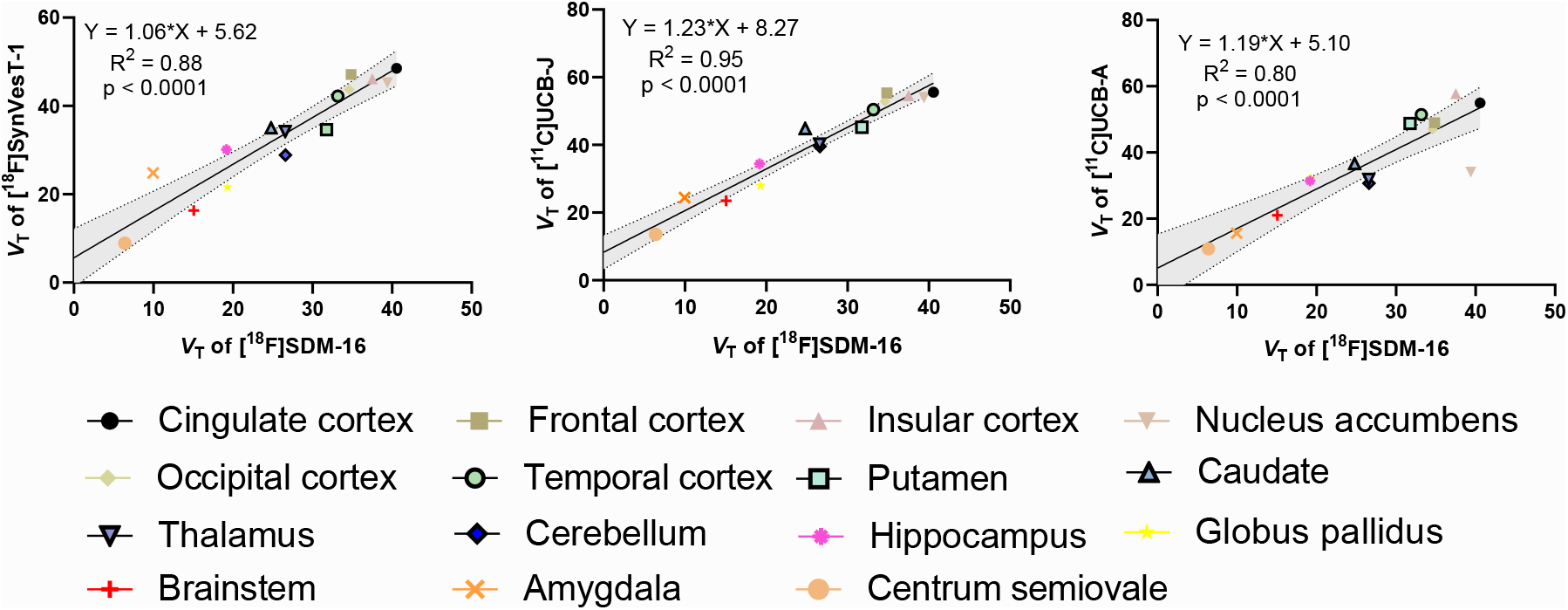
Correlation and linear regression analysis of the baseline 1TC *V*_T_ values of [^18^F]SDM-16 with that of [^18^F]SynVesT-1 [19], [^11^C]UCB-J [26], and [^11^C]UCB-A in monkey brain; dotted lines represent 95% confidence intervals.

The regional *K*_1_ values of [^18^F]SDM-16 were comparable to those of [^11^C]UCB-A, and 86% and 82% lower than those of [^18^F]SynVesT-1 and [^11^C]UCB-J, respectively (**Table 2**).

**Table 2:**
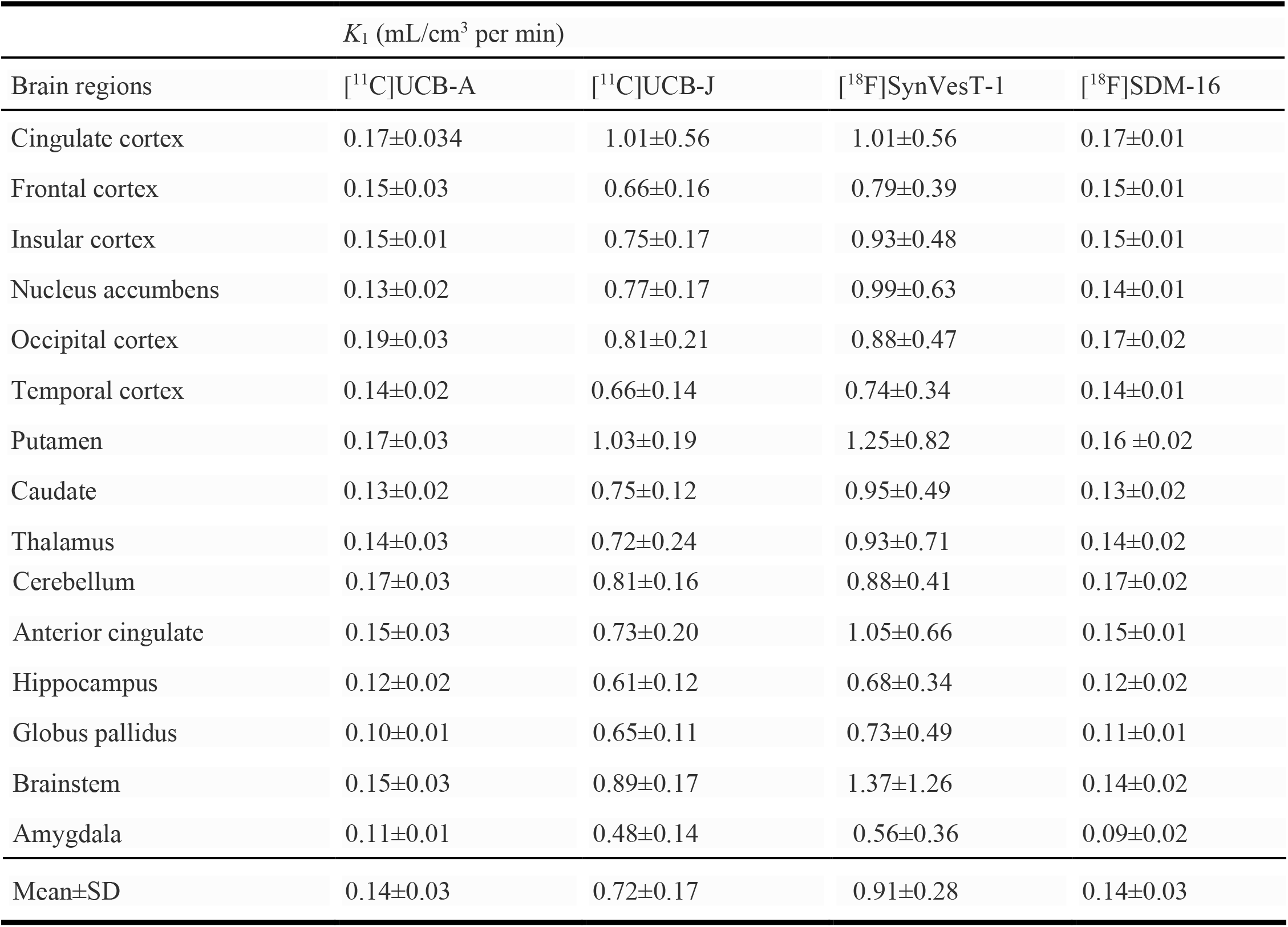
First-order kinetic rate constant (*K*_1_) of [^18^F]SDM-16 (n=4), [^18^F]SynVesT-1 (n=3) and [^11^C]UCB-J (n=5) and [^11^C]UCB-A (n=5), representing tracer influx from blood to tissue in rhesus monkey.

| Brain regions | $K_1$ (mL/cm<sup>3</sup> per min) | | | |
| --- | --- | --- | --- | --- |
| | $[^{11}\text{C}]\text{UCB-A}$ | $[^{11}\text{C}]\text{UCB-J}$ | $[^{18}\text{F}]\text{SynVesT-1}$ | $[^{18}\text{F}]\text{SDM-16}$ |
| Cingulate cortex | 0.17±0.034 | 1.01±0.56 | 1.01±0.56 | 0.17±0.01 |
| Frontal cortex | 0.15±0.03 | 0.66±0.16 | 0.79±0.39 | 0.15±0.01 |
| Insular cortex | 0.15±0.01 | 0.75±0.17 | 0.93±0.48 | 0.15±0.01 |
| Nucleus accumbens | 0.13±0.02 | 0.77±0.17 | 0.99±0.63 | 0.14±0.01 |
| Occipital cortex | 0.19±0.03 | 0.81±0.21 | 0.88±0.47 | 0.17±0.02 |
| Temporal cortex | 0.14±0.02 | 0.66±0.14 | 0.74±0.34 | 0.14±0.01 |
| Putamen | 0.17±0.03 | 1.03±0.19 | 1.25±0.82 | 0.16 ±0.02 |
| Caudate | 0.13±0.02 | 0.75±0.12 | 0.95±0.49 | 0.13±0.02 |
| Thalamus | 0.14±0.03 | 0.72±0.24 | 0.93±0.71 | 0.14±0.02 |
| Cerebellum | 0.17±0.03 | 0.81±0.16 | 0.88±0.41 | 0.17±0.02 |
| Anterior cingulate | 0.15±0.03 | 0.73±0.20 | 1.05±0.66 | 0.15±0.01 |
| Hippocampus | 0.12±0.02 | 0.61±0.12 | 0.68±0.34 | 0.12±0.02 |
| Globus pallidus | 0.10±0.01 | 0.65±0.11 | 0.73±0.49 | 0.11±0.01 |
| Brainstem | 0.15±0.03 | 0.89±0.17 | 1.37±1.26 | 0.14±0.02 |
| Amygdala | 0.11±0.01 | 0.48±0.14 | 0.56±0.36 | 0.09±0.02 |
| Mean±SD | 0.14±0.03 | 0.72±0.17 | 0.91±0.28 | 0.14±0.03 |

#### Lassen plot

To examine the in vivo binding specificity, SV2A occupancy, and the extent of nonspecific binding in the monkey brain, we performed the Lassen plot analysis using data from the two baseline scans and one blocking scan in the same monkey. The preinjected SV2A ligand LEV (30 mg/kg, i.v.) blocked 79% of the available SV2A binding sites in all grey matters (R^2^ = 0.99), indicating high *in vivo* binding specificity of [^18^F]SDM-16 (**Fig. 5**). The degree of SV2A occupancy by LEV was similar to previously reported with other SV2A PET tracers [19, 21, 26]. Based on the Lassen plot, the *V*_ND_ of [^18^F]SDM-16 in the monkey we imaged was 2.54 mL/cm^3^, which was lower than we previously determined for [^18^F]UCB-H (7.89 mL/cm^3^) [35], [^11^C]UCB-J (6.27 mL/cm^3^), [^11^C]UCB-A (14.67 mL/cm^3^), and [^18^F]SynVesT-1 (4.96 mL/cm^3^), but higher than that of [^18^F]SynVesT-2 (2.10 mL/cm^3^).

**Figure 5:**
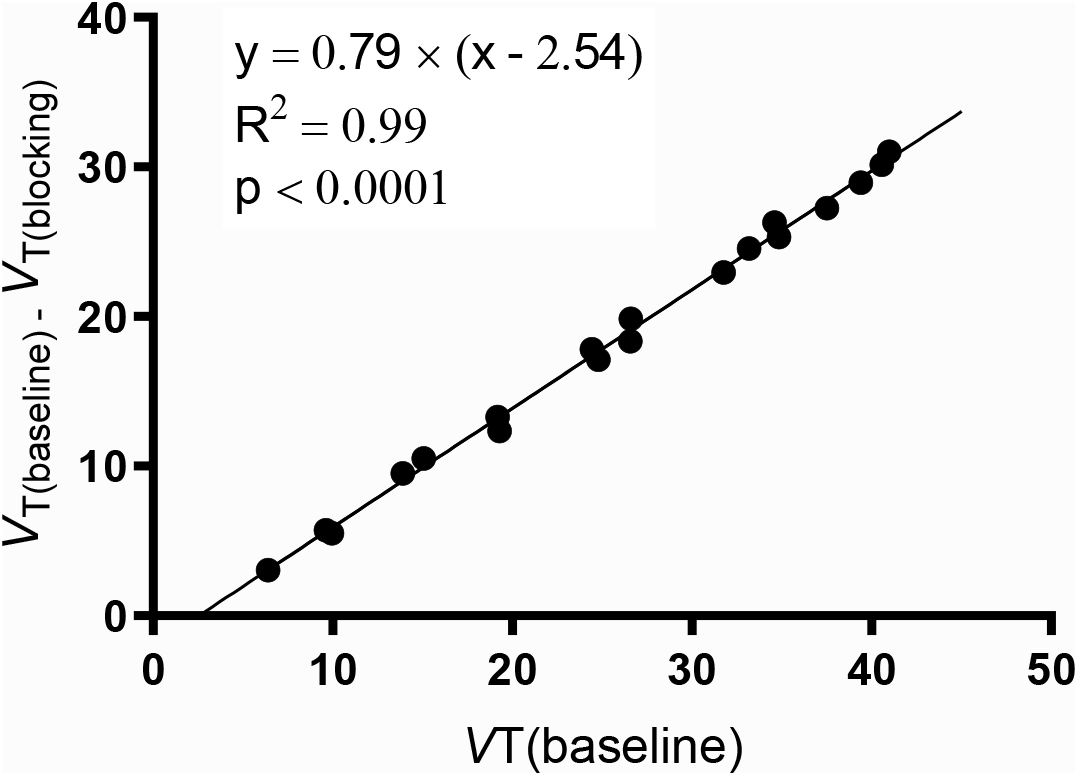
SV2A occupancy plot using the averaged *V*_T_ values from two baseline scans of [^18^F]SDM-16 and one blocking scan with preinjected LEV (30 mg/kg, i.v.) in the same rhesus monkey. The estimated SV2A occupancy by LEV (30 mg/kg, i.v.) was 79%, and the estimated nondisplaceable volume of distribution (*V*_ND_) was 2.54 mL/cm^3^.

#### Binding potential

The specific to nonspecific binding signal, as reflected by the non-displaceable binding potential (*BP*_ND_), was calculated either using the nondisplaceable volume of distribution (*V*_ND_) obtained from the blocking study or using the CS *V*_T_ (*V*_T(CS)_) as the reference. With the *V*_ND_ method, regional *BP*_ND_ values ranged from 2.9 to 15.1 (**Table 3**). Using CS as the reference region, regional *BP*_ND_ values ranged from 0.38 to 4.90, which was in average 77% lower than those calculated using *V*_ND_ values. This difference is expected due to the substantial partial volume effect in CS, resulting in an overestimation of *V*_ND_ when using CS *V*_T_. The regional *BP*_ND_ values of [^18^F]SDM-16 correlated well with those of [^18^F]SynVesT-1 (with *V*_ND_ method: Y = 0.50*X + 1.28, R^2^ = 0.84, p < 0.0001; with *V*_T(CS)_ method: Y = 0.69*X + 0.64, R^2^ = 0.80, p < 0.0001), and [^11^C]UCB-J (with *V*_ND_ method: Y = 0.42*X + 1.2, R^2^ = 0.89, p < 0.0001; with *V*_T(CS)_ method: Y = 0.54*X + 0.38, R^2^ = 0.90, p < 0.0001) (**Fig. 6**).

**Table 3:** Regional binding potentials (*BP*_ND_) of [^18^F]SDM-16 (n=2), [^18^F]SynVesT-1 (n=2) and [^11^C]UCB-J (n=5) in rhesus monkey brains.

| Brain regions | [$^{18}\text{F}$]SDM-16 | | [$^{18}\text{F}$] SynVesT-1 | | [$^{11}\text{C}$]UCB-J | |
| --- | --- | --- | --- | --- | --- | --- |
| | (mean ± SD) | | (mean ± SD) | | (mean ± SD) | |
| | $V_{ND}$ method<br>(n=2) | $V_{T(CS)}$ method<br>(n=3) | $V_{ND}$ method<br>(n=2) | $V_{T(CS)}$ method<br>(n=3) | $V_{ND}$ method<br>(n=5) | $V_{T(CS)}$ method<br>(n=5) |
| Cingulate cortex | 14.97±0.28 | 5.37±0.48 | 9.04±3.59 | 4.5±0.4 | 7.37±1.18 | 3.17±0.84 |
| Frontal cortex | 12.72±0.76 | 4.48±0.62 | 8.64±3.59 | 4.3±0.6 | 7.28±1.00 | 3.17±0.84 |
| Insular cortex | 13.77±1.93 | 4.93±1.11 | 8.41±3.27 | 4.2±0.3 | 7.24±0.73 | 3.12±0.81 |
| Nucleus accumbens | 14.51±0.23 | 5.17±0.27 | 8.41±3.54 | 4.1±0.4 | 7.16±1.13 | 3.12±0.81 |
| Occipital cortex | 12.62±1.09 | 4.45±0.75 | 7.61±2.51 | 3.9±0.2 | 7.01±0.88 | 2.97±0.65 |
| Temporal cortex | 12.06±0.54 | 4.21±0.52 | 7.36±2.64 | 3.7±0.3 | 6.57±0.89 | 2.79±0.61 |
| Putamen | 11.50±0.8 | 4.00±0.62 | 6.41±2.96 | 2.9±0.4 | 6.00±0.24 | 2.42±0.63 |
| Caudate | 8.76±0.68 | 2.90±0.50 | 6.23±2.96 | 3.0±0.4 | 5.78±0.43 | 2.40±0.72 |
| Thalamus | 9.46±0.19 | 3.16±0.17 | 6.59±3.72 | 2.9±0.7 | 5.21±0.57 | 2.03±0.62 |
| Cerebellum | 9.47±0.65 | 3.18±0.50 | 4.74±1.81 | 2.2±0.2 | 4.43±0.83 | 1.95±0.36 |
| Anterior cingulate | 15.13±4.21 | 5.33±1.30 | 8.44±4.27 | 4.1±0.7 | 6.84±0.18 | 2.70±0.74 |
| Hippocampus | 6.55±0.97 | 2.03±0.56 | 5.19±2.56 | 2.4±0.4 | 4.47±0.14 | 1.59±0.39 |
| Globus pallidus | 6.60±0.33 | 2.02±0.05 | 3.42±1.90 | 1.4±0.3 | 3.31±0.38 | 1.10±0.28 |
| Brainstem | 4.93±0.15 | 1.36±0.20 | 2.30±0.99 | 0.8±0.0 | 2.61±0.44 | 0.76±0.21 |
| Amygdala | 2.92±0.25 | 0.57±0.19 | 4.24±4.05 | 1.9±1.1 | 2.77±0.06 | 0.85±0.34 |

**Figure 6:**
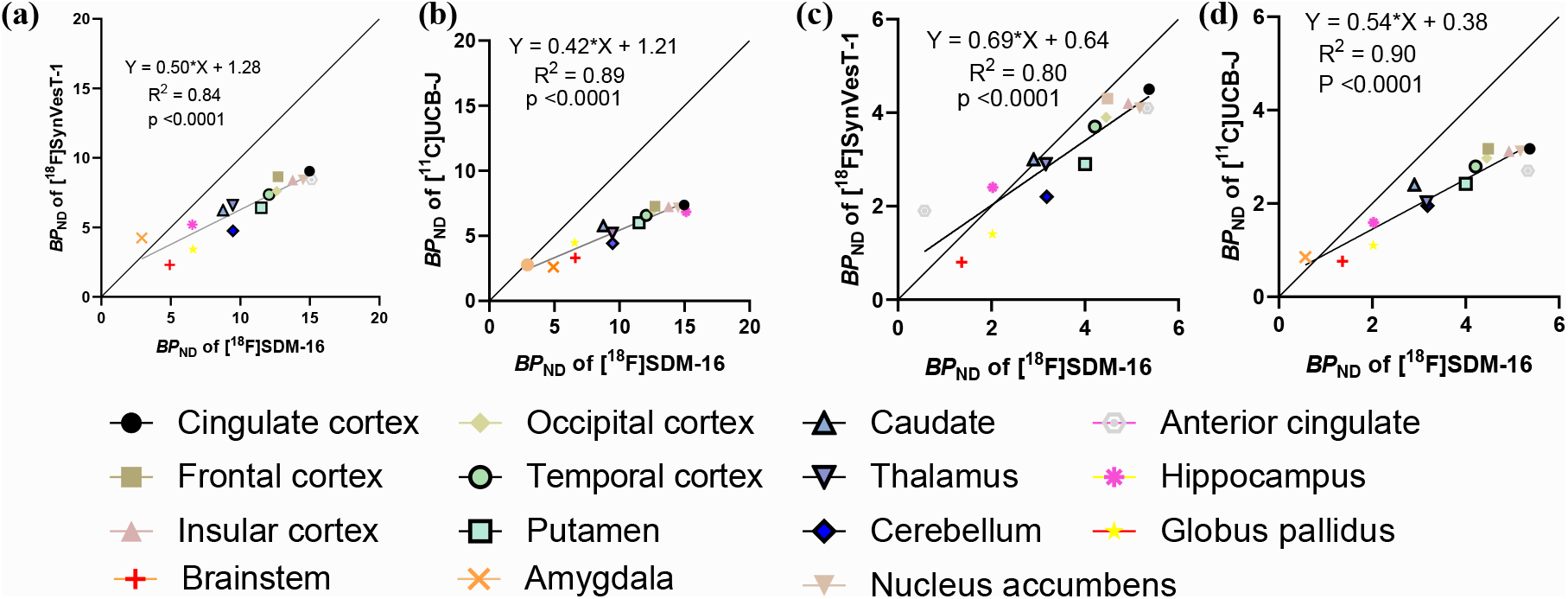
Correlation and linear regression analysis of the regional *BP_ND_* values from baseline scans of [^18^F]SDM-16 with [^18^F]SynVesT-1 and [^11^C]UCB-J, using *V*_ND_ (a, b) or centrum semiovale as reference region (c, d).

#### Test-retest reproducibility

For a preliminary evaluation of the reproducibility of the PK parameter estimation, we scanned one monkey twice with 161 days in between, using [^18^F]SDM-16. The metabolite-corrected plasma input functions and SUV TACs were highly consistent between the two scans. The 1TC *V*_T_ values of the test and retest scans correlated very well (Y = 1.09*X - 0.01, R^2^ = 0.94, P < 0.0001), with test-retest variability (TRV) for [^18^F]SDM-16 of −9.2 ± 8.5%, 11.1 ± 3.0%, and −10.4 ± 9.6% for *V*_T_, *K*_1_, and *BP*_ND_, respectively. The absolute test-retest variability (aTRV) for [^18^F]SDM-16 of 10.00 ± 6.86%, 11.14 ± 3.03%, and 11.64 ± 7.85% for *V*_T_, *K*_1_, and *BP*_ND_, respectively (**Table 4**), indicated good agreement between the two baseline scans.

**Table 4:**
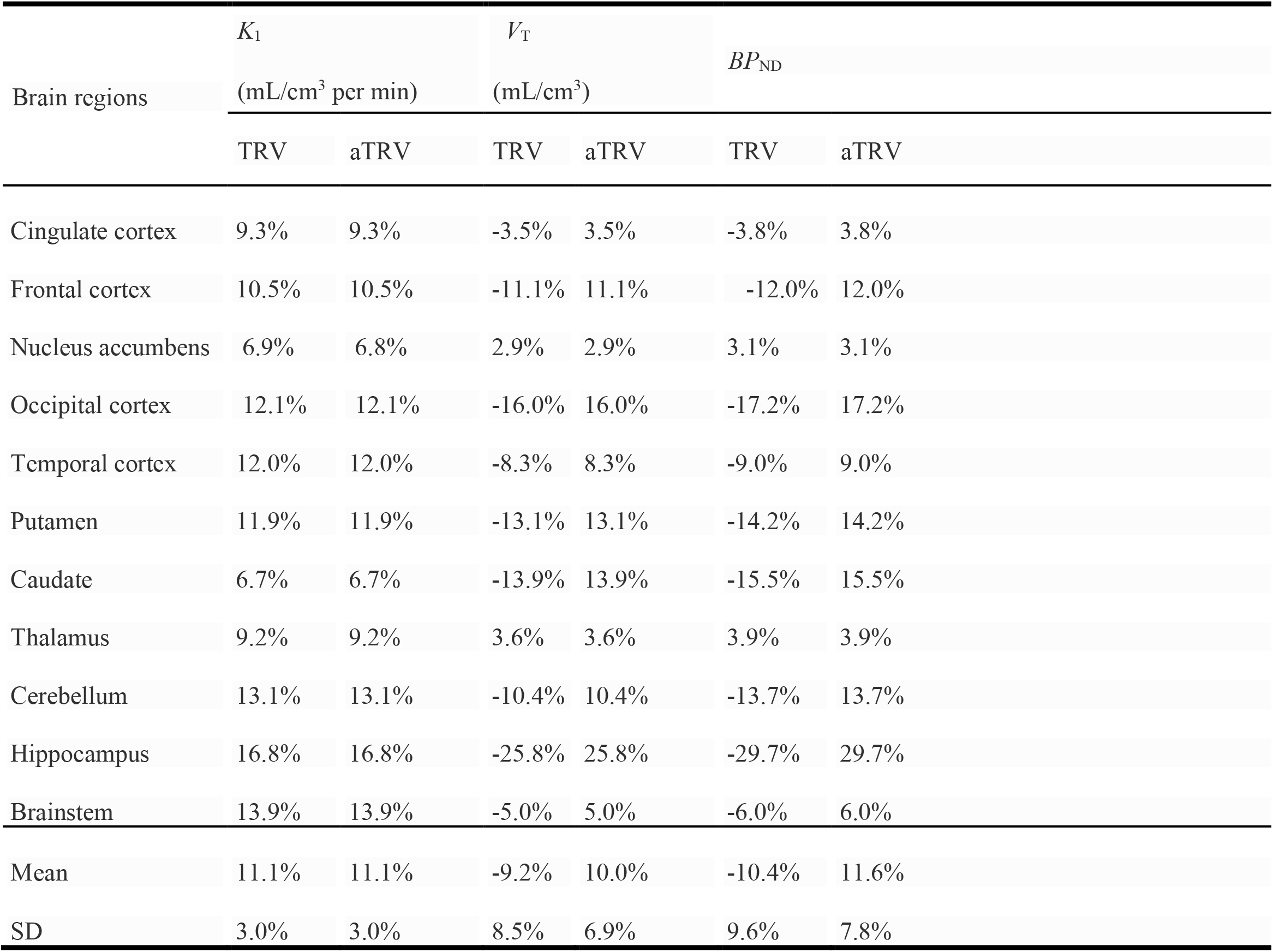
Absolute test-retest reproducibility (aTRV) of *K*_1_, *V*_t_, and *BP*_ND_ of [^18^F]SDM-16 derived with the one-tissue compartment model from 180 min PET data. *BP*_ND_ was calculated from *V*_T_ using *V*_ND_. TRV= (test value-retest value)/(test vale + retest value) × 2.

| Brain regions | $K_1$ | | $V_T$ | | $BP_{ND}$ | |
| --- | --- | --- | --- | --- | --- | --- |
|  | (mL/cm<sup>3</sup> per min) |  | (mL/cm<sup>3</sup>) |  |  |  |
|  | TRV | aTRV | TRV | aTRV | TRV | aTRV |
| Cingulate cortex | 9.3% | 9.3% | -3.5% | 3.5% | -3.8% | 3.8% |
| Frontal cortex | 10.5% | 10.5% | -11.1% | 11.1% | -12.0% | 12.0% |
| Nucleus accumbens | 6.9% | 6.8% | 2.9% | 2.9% | 3.1% | 3.1% |
| Occipital cortex | 12.1% | 12.1% | -16.0% | 16.0% | -17.2% | 17.2% |
| Temporal cortex | 12.0% | 12.0% | -8.3% | 8.3% | -9.0% | 9.0% |
| Putamen | 11.9% | 11.9% | -13.1% | 13.1% | -14.2% | 14.2% |
| Caudate | 6.7% | 6.7% | -13.9% | 13.9% | -15.5% | 15.5% |
| Thalamus | 9.2% | 9.2% | 3.6% | 3.6% | 3.9% | 3.9% |
| Cerebellum | 13.1% | 13.1% | -10.4% | 10.4% | -13.7% | 13.7% |
| Hippocampus | 16.8% | 16.8% | -25.8% | 25.8% | -29.7% | 29.7% |
| Brainstem | 13.9% | 13.9% | -5.0% | 5.0% | -6.0% | 6.0% |
| Mean | 11.1% | 11.1% | -9.2% | 10.0% | -10.4% | 11.6% |
| SD | 3.0% | 3.0% | 8.5% | 6.9% | 9.6% | 7.8% |

#### Dosimetry

In preparation for the evaluation of [^18^F]SDM-16 in humans, we performed a whole-body distribution study in one rhesus monkey. Organ residence times are shown in **Table S1**, while the absorbed doses estimated for the 55 kg reference female phantom using the OLINDA/EXM software (Vanderbilt University) were listed in **Table S2**. The organ receiving the largest dose was the urinary bladder wall (0.1368 mGy/MBq), followed by the brain (0.1032 mGy/MBq), liver (0.0538 mGy/MBq), kidneys (0.0454 mGy/MBq), and the gallbladder wall (0.0441 mGy/MBq). Based on the urinary bladder wall as the critical organ, the maximum permissible single study dosage of [^18^F]SDM-16, to remain below the 21 CFR 361.1 dose limit, is 365.6 MBq (9.88 mCi). The estimated effective dose (ED) is 21.1 μSv/MBq, is slightly higher than the reported value of 15.4 μSv/MBq estimated for [^18^F]UCB-H from human [18], and is similar to the ED value of 20 μSv/MBq for [^18^F]SynVesT-1 estimated from female rhesus macaques using the 1 hour voiding model [37], and is within the range of ED values (15 – 29 μSv/MBq) reported for [^18^F]FDG [38–41]. The total ED resulting from a single study dosage of 185 MBq (5 mCi) [^18^F]SDM-16 is estimated to be equivalent to 3.9 mSv (0.39 rem). Accordingly multiple PET scans can be performed within the same research subject based on individual’s whole body annual and total dose commitment of 50 mSv (21 CFR 361.1).

## DISCUSSION

A quantitative tool to image the CNS-PNS synaptic axis will open the opportunity to study the interplay between brain, spinal cord and peripheral nervous system, under normal and disease conditions. We have previously reported the synthesis and evaluation of a series of fluorine-18-labeled SV2A PET tracers, which all showed excellent brain imaging properties, and some have been translated into first-in-human studies [37, 42, 43]. However, these PET tracers are metabolically labile, with less than 50% parent fraction at 30 min post injection. While our data indicate that the radiometabolites are not brain penetrant and would not interfere with the quantitative analysis of brain SV2A expression levels, these radiotracers are not suitable for imaging of SV2A in peripheral organs. For [^11^C]UCB-J and [^18^F]UCB-H, the prevalent radiometabolites in plasma are the corresponding *N*-oxidation products, which do not enter the brain to a significant extent as reported in LC/MS/MS and small animal PET imaging studies [26] [27]. Thus, we designed a new SV2A radiotracer, based on the structure of UCB-A, which possesses an imidazole ring and lacks the formation of a pyridinyl *N*-oxide radiometabolite [44, 45]. We modified the structure of UCB-A, in a way to fine-tune the physicochemical properties and further improve its in vivo stability and brain kinetics, because UCB-A’s PK in human brain is too slow to allow for the reliable estimation of PK parameters using data from a C-11 PET scan with reasonable length [46]. Based on the ChemDraw (Version 20.1.0.112)-predicted LogP values of SDM-16 (2.06) and UCB-A (0.96), we expected to see higher membrane permeability of SDM-16 over UCB-A, as in general, within the same series of compounds, higher lipophilicity is associated with higher cell membrane permeability [47]. However, since a higher fraction of SDM-16 is expected to be protonated at physiological pH than UCB-A, the delivery of SDM-16 from plasma to brain could potentially be hampered if the positively charged molecule does not enter the brain as effectively as the uncharged molecule.

The newly designed SV2A ligand SDM-16 binds to human SV2A with high affinity as a racemic mixture. Based on our experience with the synthesis of [^18^F]SynVesT-1 [48] and [^18^F]SynVest-2 using organotin precursors [21], we decided to apply the same radiolabeling strategy for [^18^F]SDM-16. To our satisfaction, [^18^F]SDM-16 was synthesized with high radiochemical yield, radiochemical and chemical purities, and molar activities. The relatively higher hydrophilicity of SDM-16 than UCB-J, SynVesT-1, and SynVesT-2 was expected to increase its free fractions in plasma and brain. Indeed, the plasma free fraction (*f*_P_) of [^18^F]SDM-16 is 69%, which is slightly lower than that of [^11^C]UCB-A (75%), but much higher than of [^11^C]UCB-J (46%), [^18^F]SynVesT-1 (43%), [^18^F]SynVesT-2 (41%), and [^18^F]UCB-H (43%) (**Fig. 7**). The trend in *f*_P_ is consistent with the relative measured lipophilicity of [^11^C]UCB-J (LogP: 2.46), [^18^F]SynVesT-1 (LogP: 2.32), [^18^F]SynVesT-2 (LogP: 2.17), [^18^F]SDM-16 (LogP: 1.65), and [^11^C]UCB-A (LogP: 1.10), with *f*_P_ negatively correlated with LogP (R^2^ = 0.87, P = 0.02). The clearance rate of [^18^F]SDM-16 from plasma is slower than [^18^F]SynVesT-1, [^11^C]UCB-J and [^11^C]UCB-A, and we observed the highest parent fractions for [^18^F]SDM-16 among all the SV2A PET tracers we evaluated in monkey. The relatively slower plasma clearance (**Fig. 2a**) and higher plasma parent fraction (**Fig. 2b**) indicate the higher metabolic stability of [^18^F]SDM-16 over the other existing SV2A radiotracers.

**Figure 7:**
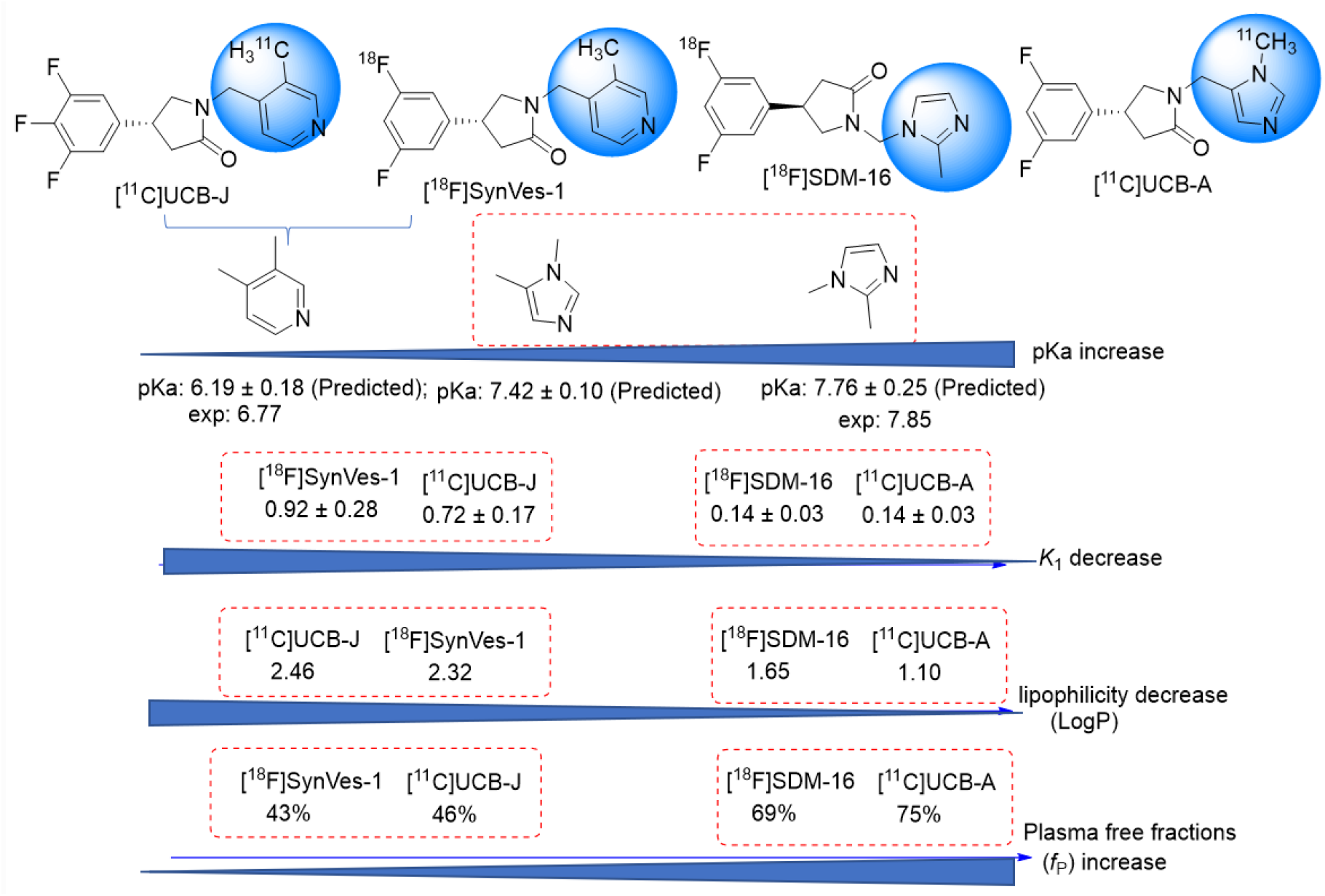
The influence of pK_a_ on the properties of PET tracers.

Because of the high metabolic stability and consistently high tracer concentration in the plasma (**Fig. 2a**), the brain TACs of [^18^F]SDM-16 appear similar to tracers with irreversible binding kinetics (**Fig. 3c**). As for other SV2A PET tracers, the 1TC model provided good fits and reliable estimates of PK parameters of [^18^F]SDM-16 (**Fig. 3c**). The excellent 1TC fitting and the efficient displacement by LEV (**Fig. 3e**) demonstrate the reversible binding kinetics of [^18^F]SDM-16. Thus, we obtained the *K*_1_ and *V*_T_ parameters using the 1TC model. The *K*_1_ values of [^18^F]SDM-16 are comparable to those of [^11^C]UCB-A and are much lower than those of [^18^F]SynVesT-1 and [^11^C]UCB-J, respectively (**Table 2**). There are many factors that could influence the delivery of drug molecules from plasma into the brain, e.g., passive membrane permeability, plasma free fraction, ionization state in the plasma or cytosol, active transportation and efflux, etc. Based on the topological analysis of the structures of the current SV2A PET tracers, both [^18^F]SDM-16 and [^11^C]UCB-A are imidazole derivatives; while [^18^F]SynVesT-1, [^18^F]SynVesT-2, and [^11^C]UCB-J share a common lutidine substructure. Considering their common fluorophenylpyrrolidin-2-one pharmacophore, which is unconjugated with the pyridine/imidazole, their acid/base properties are mainly driven by the different heteroaromatic substituents. We speculate that the ionization constants (pKa values) of the conjugate acids of these four SV2A ligands affects their *K*_1_ values. According to the calculations using the Advanced Chemistry Development Software (ACD/Labs, V11.02) and reported experimental data [49-51], *N*-protonated lutidine has higher pKa than *N*-protonated dimethyl imidazole (**Fig. 7**). Modification of the Henderson-Hasselbalch equation leads to the equation to calculate the concentration ratio of the tracer in free base form [B] to that of the total 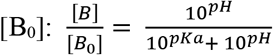 (**Eq. 1**). According to **Eq. 1**, 81% of [^18^F]SynVesT-1, 49% of [^11^C]UCB-A, and 30% of [^18^F]SDM-16 would be present in the plasma as free bases, based on the experimental pKa data of lutidine (6.77) and dimethyl imidazole (7.42, 7.85), and the pH value of plasma being 7.4.

The acid/base property is a critical factor to consider in drug discovery, especially for CNS drugs that have special requirement for BBB penetration [52]. The pKa values of all these SV2A ligands are within the commonly accepted range for pKa of CNS drugs, i.e., from 4 to 10 [34]. While the higher plasma free fraction of SDM-16 favors its delivery from plasma into the brain, the more extensive protonation of SDM-16 and its relatively higher polar surface area could contribute to its lower *K*_1_ values than [^11^C]UCB-J and [^18^F]SynVesT-1. Considering the nearly identical *K*_1_ values of [^18^F]SDM-16 and [^11^C]UCB-A (Y = 0.97*X + 0.01, R^2^ = 0.86, P < 0.0001) and that the averaged *V*_T_ of [^18^F]SDM-16 was 71% of that of [^11^C]UCB-A, [^18^F]SDM-16 is expected to reach brain-to-blood distribution equilibrium 1.4-fold faster than [^11^C]UCB-A. To estimate the time for [^18^F]SDM-16 to reach brain equilibration, we calculated the equilibration half-lives for [^18^F]SDM-16 and [^11^C]UCB-A in the selected brain regions (*t*_1/2_ = ln2/k_2_, **Table S3**). The brain equilibration half-life of [^18^F]SDM-16 is about the half-life of ^18^F, ranging from 84 ± 9 min in CS to 201 ± 54 min in insular cortex (**Table S3**). In average, the brain equilibration half-life of [^11^C]UCB-A is about 1.4 fold longer than that of [^18^F]SDM-16. Although [^18^F]SDM-16 displaced relatively slow kinetics in the rhesus monkey brain where SV2A expression level is high (SV2A *B*_max_ of Baboon’ brain ranged from 2.2 pmol/mg protein in the pons to 19.9 pmol/mg protein in the temporal cortex) [5], its kinetics is expected to be faster in tissues with relatively lower SV2A expression, such as spinal cord [53].

Although in the field of PET neuroimaging, the rule of thumb is that the *B*_max_/*K*_d_ of the PET tracer needs to be greater than 10, the ratio of two tracers’ *BP*_ND_ is determined partially by their degree of nonspecific uptake, which is reflected in *V*_ND_ and brain tissue free fraction (*f*_ND_). While *V*_ND_ can be obtained only through in vivo blocking studies, *f*_ND_ can be obtained either from in vivo blocking study or from in vitro assays using brain homogenates or slides, and *f*_ND_ is considered to be consistent among different species [54]. While decreasing the tracer’s *K*_d_ value may eventually leads to undesired slow kinetics (low *k*_2_ and long brain-to-plasma equilibrium half-life), increasing *f*_ND_ is an alternative but potentially more challenging approach to boost the specific PET signal, based on the equation *BP*_ND_ = *f*_ND_\**B*_max_/*K*_d_. Using the averaged *V*_ND_ and *f*_P_ values, we calculated the *f*_ND_ value of [^18^F]SDM-16 to be 27%, which was higher than that of [^18^F]UCB-H (6.1%, calculated from *K*_1_/k_2_ using 2TCM-c) [35], [^11^C]UCB-J (7.3%), [^18^F]SynVesT-1 (10.1%), and [^18^F]SynVesT-2 (19.5%), assuming that these SV2A ligands enter the brain mainly through passive diffusion and are not subject to active influx or efflux transport, i.e., *f*_ND_ = *f*_P_/*V*_ND_. We did not calculate the *V*_ND_ and *f*_ND_ values of [^11^C]UCB-A because of the lack of blocking data for [^11^C]UCB-A. [^18^F]SDM-16 has the highest *f*_ND_ value among all the current SV2A PET tracers, and maintains the brain penetration ability.

To compare the *in vivo K*_d_ and *BP*_ND_ of [^18^F]SDM-16 with those of [^18^F]SynVesT-1 [19], [^11^C]UCB-J [26], and [^11^C]UCB-A in monkey brain, we adopted the Guo plot using their baseline *V*_T_ values (**Fig. 4**). The *K*_d_ ratios are *K*_d_([^18^F]SynVesT-1)/*K*_d_([^18^F]SDM-16)=0.52 and *K*_d_([^11^C]UCB-J)/*K*_d_([^18^F]SDM-16)=0.51; while the y-intercepts are greater than zero, indicating higher *BP*_ND_ of [^18^F]SDM-16 than [^18^F]SynVesT-1, [^11^C]UCB-J and [^11^C]UCB-A. The *BP*_ND_ ratios are *BP*_ND_([^18^F]SDM-16)/*BP*_ND_([^18^F]SynVesT-1)=1.60 and *BP*_ND_([^18^F]SDM-16)/*BP*_ND_([^11^C]UCB-J)=2.94. Because we used different monkeys in the evaluations of these SV2A PET tracers, the *BP*_ND_ ratios or *in vivo K*_d_ ratios could be influenced by animal differences.

Next, we calculated the *BP*_ND_ values of the SV2A PET tracers using either CS as reference region or using the *V*_ND_ derived from blocking studies. We noticed that the *BP*_ND_ values calculated using the *V*_T_ values of CS are 67.2 ± 4.4% lower than the true *BP*_ND_ derived from *V*_ND_ values. Contributing factors to the underestimation of *BP*_ND_ using CS as reference region are the spill-in effect of the PET signal from the gray matter surrounding CS and the presence of SV2A specific uptake in CS. The ranking order of [^18^F]SDM-16 *BP*_ND_ values in all ROIs (cingulate cortex > frontal cortex > insula > temporal cortex > putamen > caudate > cerebellum > hippocampus > brainstem > amygdala) is basically consistent with those of [^11^C]UCB-A, [^18^F]SynVesT-1, and [^11^C]UCB-J (**Table 3**). Note that the *BP*_ND_ values of [^18^F]SDM-16 calculated using both methods are generally higher than those of [^11^C]UCB-A, [^18^F]SynVesT-1, and [^11^C]UCB-J (**Table 3** and **Fig. 6**), which is consistent with the Guo plot analysis results. However, since the monkeys used in each tracer’s evaluation are different, further studies using the same cohort of monkeys are needed to confirm if [^18^F]SDM-16 possess higher specific binding than the other SV2A PET tracers in NHP brains. An SV2A PET tracer with high specific binding signals will be advantageous in the imaging and quantification of SV2A in tissues with relatively low SV2A expression, e.g., spinal cord [53] and pancreas [55]. In fact, the *BP*_ND_ values of [^18^F]SDM-16 in the LEV blocking scan are relatively high in the gray matters (up to 2.11 in cingulate cortex and nucleus accumbens), even with 79% of the SV2A being occupied by LEV, indicating that [^18^F]SDM-16 is advantageous in the imaging and quantification of SV2A at much less densities than the cerebrum.

## CONCLUSIONS

We have successfully synthesized a new ^18^F-labeled SV2A PET tracer [^18^F]SDM-16 and evaluated its imaging characteristics in rhesus monkeys. [^18^F]SDM-16 is metabolically more stable than the current SV2A PET tracers, displayed reversible and high specific binding in NHP brain with relatively low nonspecific binding in white matter. The TACs fitted well with 1TC to allow for reliable estimation of PK parameters. [^18^F]SDM-16 may have potential applications in the quantification of SV2A in the whole CNS and possibly in the PNS and neuroendocrine system as well.

## ACKNOWLEDGEMENT

The authors thank the professional technical support by the Yale PET Center staff. This study was supported by grants from the National Institutes of Health (NIH) and the Archer Foundation. The contents are solely the responsibility of the authors and do not necessarily represent the official view of the funding agencies.

