## Supporting Information for "A metabolically stable PET tracer for imaging synaptic vesicle protein 2A: Synthesis and preclinical characterization of [^18^F]SDM-16"

### Methods

**Chemistry and general conditons**. All reagents and solvents were purchased from commercial sources (Sigma-Aldrich, VWR, and Fisher Scientific, etc.) and used without further purification. Proton, carbon, and fluorine nuclear magnetic resonance ($^1$H, $^{13}$C, and $^{19}$F NMR) spectra were recorded on an Agilent 400 or 500 MHz spectrometer. Chemical shifts are reported in parts per million (ppm), with either the solvent resonance or tetramethyl silane (TMS) as the internal standard (TMS, 0 ppm; $CDCl_3$, 7.26 ppm; DMSO-$d_6$, 2.49 ppm in $^1$H NMR and $CDCl_3$, 77.0 ppm; DMSO-$d_6$, 39.7 ppm in $^{13}$C NMR). Multiplicities are indicated as s (singlet), d (doublet), t (triplet), q (quartet), quint (quintet), m (multiplet); coupling constants *J* are given in hertz (Hz). All chemicals used in this study were of ≥ 95% purity based on HPLC (detection at 254 nm), or NMR. HRMS (ESI) data were tested on a Shimadzu 9030 QToF LC-MS system.

**Chiral Separation of 4-(3,5-difluorophenyl)pyrrolidin-2-one (9)**. Compound **9** was synthesized in 45% yield over 3 steps according to the previously reported procedure [1]. The separation was conducted on a ChiralPak AS-H preparative HPLC column (250 x 30 mm I.D., 5µm), eluting with n-heptane/EtOH (3 : 2 v/v), with the active enantiomer (*R*)-**9** eluting first, followed by the inactive enantiomer (*S*)-**9**. The *e.e.* was determined on an analytical Chiralpak IA column (5 µm, 4.6 x 150 mm) eluting with hexane/EtOH (1 : 1 v/v) at a flow rate of 1.00 mL/min. Both enantiomers showed >99% ee. (*R*)-**9**: $R_t$ = 5.81 min, (*S*)-**9**: $R_t$ = 6.25 min. $^1$H NMR ($CDCl_3$, 400 MHz): $\delta$ 6.78 (d, *J* = 6.44 Hz, 2H), 6.77-6.63 (m, 1H), 6.61 (s, 1H), 3.78 (t, *J* = 10.0 Hz, 1H), 3.65 (q, *J* = 16.0 Hz, 1H), 3.38 (t, *J* = 8.0 Hz, 1H), 2.74 (dd, *J* = 16.0, 8.0 Hz, 1H), 2.42 (dd, *J* = 16.0, 8.0 Hz, 1H). $^{13}$C NMR ($CDCl_3$, 100 MHz): $\delta$ 37.5, 39.8, 48.9, 102.6 (t, *J* = 25.0 Hz), 109.7 (dd, *J* = 19.0, 7.0 Hz, 2C), 146.0 (t, *J* = 9.0 Hz), 163.2 (dd, *J* = 248.0, 13.0 Hz, 2C), 176.9. $^{19}$F NMR ($CDCl_3$, 376 MHz): $\delta$ -108.96.

**4-(3,5-difluorophenyl)-1-(hydroxymethyl)pyrrolidin-2-one (10)**. In a 2.5 ml V-vial were placed compound **9** (0.05 g, 0.253 mmol), 37% formalin solution (38 µl, 0.507 mmol) and water (0.5 ml) mixed, and heated at 115 °C for 16 h. After cooling to room temperature, the suspension was dissolved in methanol (10 ml), the solvents were removed under reduced pressure, and the residue was purified by column chromatography ($SiO_2$, 19/1/0.01 $CH_2Cl_2$/MeOH/$NH_3OH$) to give **10** (55 mg, 96%). $^1$H NMR ($CDCl_3$, 500 MHz): $\delta$ 2.50 (dd, *J* = 17.1, 7.7 Hz, 1H), 2.84 (dd, *J* = 17.1, 8.8 Hz, 1H), 3.55-3.61 (m, 2H), 3.93 (t, *J* = 8.2 Hz, 1H), 4.46 (s, 1H), 4.77 (d, *J* = 8.7 Hz, 1H), 4.91 (d, *J* = 9.2 Hz, 1H), 6.69 (t, *J* = 8.9 Hz, 1H), 6.77 (d, *J* = 5.9 Hz, 2H); $^{13}$C NMR ($CDCl_3$, 125 MHz): $\delta$ 37.0, 39.1, 52.7, 66.5, 102.8 (t, *J* = 25.3 Hz), 109.9 (dd, *J* = 19.2, 5.8 Hz, 2C), 146.3 (t, *J* = 8.8 Hz), 163.4 (dd, *J* = 249.2, 12.9 Hz, 2C), 174.4. $^{19}$F NMR ($CDCl_3$, 470 MHz): $\delta$ -108.85.

**4-(3,5-difluorophenyl)-1-((2-methyl-1*H*-imidazol-1-yl)methyl)pyrrolidin-2-one (*rac* SDM-16, 13)**. In a two-neck round bottom flask 2-methylimidazole (50 mg, 0.616 mmol), and *N*, *N*-diisopropylethylamine (100 µl, 0.572 mmol) were dissolved in acetonitrile (4.5 ml). To this mixture oxalyl chloride (22 µl, 0.264 mmol) dissolved in acetonitrile (0.5 ml) were slowly added. Under argon atmosphere the mixture was stirred for 30 min at room temperature. Then (100 mg, 0.440 mmol) ***rac*** **10** were added, stirred for 30 min at room temperature and further heated under refluxed for 5.5 h. After cooling to room temperature, the solvent was removed under reduced pressure and the crude mixture was purified by column chromatography (I: $SiO_2$, 19/1/0.01 $CH_2Cl_2$/MeOH/ $NH_3OH$ and II: $SiO_2$, gradient system 100-80% ethyl acetate/ethanol) to give **13** (59 mg, 46%). $^1$H NMR ($CDCl_3$, 400 MHz): $\delta$ 2.43 (s, 3H), 2.51 (dd, *J* = 17.2, 8.2 Hz, 1H), 2.83 (dd, *J* = 17.2, 9.0 Hz, 1H), 3.29 (dd, *J* = 9.3, 6.9 Hz, 1H), 3.49-3.57 (m, 1H), 3.69 (t, *J* = 8.8 Hz, 1H), 5.31-5.42 (m, 2H), 6.64-6.71 (m, 3H), 6.91 (s, 2H); $^{13}$C NMR ($CDCl_3$, 100 MHz): $\delta$ 12.9,

29.6, 36.8 (t, $J$ = 1.8 Hz), 37.9, 52.1 (d, $J$ = 3.3 Hz), 102.9 (t, $J$ = 25.2 Hz), 109.7 (dd, $J$ = 17.9, 7.0 Hz, 2C), 119.4, 128.1, 144.9, 145.1 (t, $J$ = 8.8 Hz), 163.3 (dd, $J$ = 249.7 Hz, $J$ = 12.9 Hz, 2C), 173.1. $^{19}$F NMR ($CDCl_3$, 376 MHz): $\delta$ -108.41. HRMS (ESI) Calcd for $C_{15}H_{16}N_3OF_2^+$ $[M+H]^+$: 292.1256, found 292,1280.

**(2-methyl-1*H*-imidazol-1-yl)methanol (11)**. 2-methyl-1*H*-imidazole (3.0 g, 36.54 mmol), paraformaldehyde (1.21 g, 40.19mmol), and triethylamine (0.037 g, 0.36 mmol) were mixed, heated at 100 $^o$C until 2-methyl-1*H*-imidazole disappear, about 30 min, afford colorless solid 4.07g. After washed with $Et_2O$ afford **11** in quantitative yield. $^1$H NMR (DMSO-$d_6$, 400 MHz): $\delta$ 2.27 (s, 3H), 5.17 (s, 2H), 6.67 (s, 1H), 7.02 (s, 1H); $^{13}$C NMR (DMSO-$d_6$, 100 MHz): $\delta$ 12.8, 68.5, 119.9, 126.4, 144.2.

**1-(chloromethyl)-2-methyl-1*H*-imidazole hydrogen chloride (12)**. Compound **11** (4.1 g, 36.54 mmol) was dissolved in thionyl chloride ($SOCl_2$) (10 ml) at 0 $^o$C, then stirred at room temperature for 3 h, the solvents were removed under reduced pressure, dried in vacuo to afforded **12** (5.5 g, 90%), mixed with 10% 2-methyl-1*H*-imidazol-1-yl)methanol hydrochloride (hydrochloride of **11**). $^1$H NMR (DMSO-$d_6$, 400 MHz): $\delta$ 2.64 (s, 3H), 6.12 (s, 2H), 7.53 (d, $J$ = 4.0 Hz, 1H), 7.75 (d, $J$ = 4.0 Hz, 1H); $^{13}$C NMR (DMSO-$d_6$, 100 MHz): $\delta$ 10.6, 53.3, 119.1, 122.4, 146.4.

**Chiral Separation of 4-(3-bromo-5-fluorophenyl)pyrrolidin-2-one (15)**. Compound **15** was synthesized in 60% yield over 3 steps according to the previously reported procedure [1]. The separation was conducted on a ChiralPak AS-H preparative HPLC column (250 x 30 mm I.D., 5µm), eluting with n-heptane/EtOH (3 : 2 v/v), with the active enantiomer (*R*)-15 eluting first, followed by the inactive enantiomer (*S*)-**15**. The *e.e.* was determined on an analytical Chiralpak IA column (5 µm, 4.6 x 150 mm) eluting with 50:50 EtOH/hexane at a flow rate of 1.00 mL/min. Both enantiomers showed >99% ee. (*R*)-**15**: $R_t$ = 5.39 min, (*S*)-**15**: $R_t$ = 5.73 min. $^1$H NMR ($CDCl_3$, 400 MHz): $\delta$ 7.17-7.11 (m, 2H), 6.88 (d, $J$ = 8.0 Hz, 1H), 6.55 (s, 1H), 3.77 (t, $J$ = 8.0 Hz, 1H), 3.62 (q, $J$ = 8.0 Hz, 1H), 3.37 (t, $J$ = 8.0 Hz, 1H), 2.71 (dd, $J$ = 16.0, 8.0 Hz, 1H), 2.41 (dd, $J$ = 16.0, 8.0 Hz, 1H). $^{13}$C NMR ($CDCl_3$, 100 MHz): $\delta$ 37.6, 39.6, 49.0, 112.8 (d, $J$ = 22.0 Hz), 117.9 (d, $J$ = 22.0 Hz), 123.0 (d, $J$ = 10.0 Hz), 125.8 (d, $J$ = 3.0 Hz), 146.2 (dd, $J$ = 8.0 Hz, 3.0 Hz), 162.5 (d, $J$ = 250.0 Hz), 177.0. $^{19}$F NMR ($CDCl_3$, 376 MHz): $\delta$ -109.83.

**(*R*)-4-(3,5-difluorophenyl)-1-((2-methyl-1*H*-imidazol-1-yl)methyl)pyrrolidin-2-one (SDM-16, *R*-13)**. To a solution of compound **(*R*)-9** (10 mg, 0.05 mmol) in anhydrous THF (0.5 ml) under argon and cooled to 0 $^o$C was added sodium hydride (NaH, 5 mg, 0.11 mmol). Tetrabutylammonium iodide (TBAI, 1 mg, 0.003 mmol) and compound **12** (20 mg, 0.12 mmol) were added after 30 min. The reaction mixture was kept stirring for 16 h at room temperature, then quenched with saturated $NaHCO_3$ solution (1 ml) and extracted with EtOAc (5 ml x 3). The combined organic phase was dried over $Na_2SO_4$ and concentrated *in vacuo*. The crude product was purified on a silica gel column eluting with 0-10% EtOH/EtOAc to afford compound **13** as an oil (13 mg, 92%). $^1$H NMR ($CDCl_3$, 400 MHz): $\delta$ $^1$H NMR ($CDCl_3$, 400 MHz): $\delta$ 2.43 (s, 3H), 2.51 (dd, $J$ = 17.2 Hz, 8.2 Hz, 1H), 2.83 (dd, $J$ = 17.2 Hz, 9.0 Hz, 1H), 3.29 (dd, $J$ = 9.3 Hz, 6.9 Hz, 1H), 3.49-3.57 (m, 1H), 3.69 (t, $J$ = 8.8 Hz, 1H), 5.31-5.42 (m, 2H), 6.64-6.71 (m, 3H), 6.91 (s, 2H); $^{13}$C NMR ($CDCl_3$, 100 MHz): $\delta$ 12.9, 29.6, 36.8 (t, $J$ = 1.8 Hz), 37.9, 52.1 (d, $J$ = 3.3 Hz), 102.9 (t, $J$ = 25.2 Hz), 109.7 (dd, $J$ = 17.9, 7.0 Hz, 2C), 119.4, 128.1, 144.9, 145.1 (t, $J$ = 8.8 Hz), 163.3 (dd, $J$ = 249.7, 12.9 Hz, 2C), 173.1. $^{19}$F NMR ($CDCl_3$, 376 MHz): $\delta$ -108.40.

**(*R*)-4-(3-bromo-5-fluorophenyl)-1-((2-methyl-1*H*-imidazol-1-yl)methyl)pyrrolidin-2-one (16)**. Compound **16** was prepared in procedures similar to those described in supporting information for ***rac* SDM-16 (13)**. Yield 90%. $^1$H NMR ($CDCl_3$, 400 MHz): $\delta$ 2.37 (s, 3H), 2.45 (dd, $J$ = 20.0, 8.0 Hz, 1H), 2.76 (dd, $J$ = 20.0, 8.0 Hz, 1H), 3.24 (t, $J$ = 8.0 Hz, 1H), 3.43-3.51 (m,

1H), 3.65 (t, $J$ = 10.0 Hz, 1H), 5.26-5.36 (m, 2H), 6.73 (d, $J$ = 12.0 Hz, 1H), 6.85 (d, $J$ = 8.0, 2H), 7.03 (s, 1H), 7.06 (d, $J$ = 8.0, 1H). $^{13}$C NMR ($CDCl_3$, 100 MHz): $\delta$ 12.9, 36.5, 36.5, 37.9, 52.0 (d, $J$ = 5.0 Hz), 112.7 (d, $J$ = 22.0 Hz), 118.1 (d, $J$ = 24.0 Hz), 119.4, 123.1 (d, $J$ = 10.0 Hz), 125.6 (d, $J$ = 3.0 Hz), 128.0, 144.8, 145.3 (d, $J$ = 8.0 Hz), 163.5 (d, $J$ = 245.0 Hz), 173.0. $^{19}$F NMR ($CDCl_3$, 376 MHz): $\delta$ -109.46.

**(*R*)-4-(3-fluoro-5-(trimethylstannyl)phenyl)-1-((2-methyl-1*H*-imidazol-1-yl)methyl)pyrrolidin-2-one (17).** To a solution of compound **16** (60 mg, 0.17 mmol) in anhydrous toluene (0.62 ml) was added lithium chloride (44 mg, 1.02 mmol), tetrakis(triphenylphosphine)palladium (20 mg, 0.02 mmol), triphenylphosphine (2 mg, 0.01 mmol), and hexamethylditin (57 µL, 90 mg, 0.27 mmol) under argon. The reaction mixture was degassed and refilled with argon for 3 min and kept stirring at 100 $^{o}$C for 1 h. The reaction mixture was diluted with EtOAc (2 ml), passed through celite, and rinsed with EtOAc (2 mL x 2). The filtrate was concentrated in vacuo. The crude product was purified on a silica gel column eluting with 0-20% EtOH/EtOAc to afford the product **17** as an oil (34 mg, 46%). $^{1}$H NMR ($CDCl_3$, 400 MHz): $\delta$ 0.20-0.33 (m, 9H), 2.44 (s, 3H), 2.56 (dd, $J$ = 16.0 Hz, 8.0 Hz, 1H), 2.83 (dd, $J$ = 16.0 Hz, 8.0 Hz, 1H), 3.30 (t, $J$ = 8.0 Hz, 1H), 3.54 (m, 1H), 3.68 (t, $J$ = 8.0 Hz, 1H), 5.30-5.43 (m, 2H), 6.74-6.77 (m, 1H), 6.90 (s, 2 H), 6.99 (s, 1H), 7.04-7.06 (m, 1H); $^{13}$C NMR ($CDCl_3$, 100 MHz): $\delta$ -9.4 (3H), 12.9, 36.9, 38.2, 52.1, 52.6, 113.2 (d, $J$ = 22.0 Hz), 119.5, 121.1 (d, $J$ = 17.00 Hz), 121.5 (d, $J$ = 2.00 Hz), 143.1 (d, $J$ = 6.00 Hz), 144.9, 146.5, 162 (d, $J$ = 251 Hz), 164.0, 173.7.

1. Li S, Cai Z, Wu X, Holden D, Pracitto R, Kapinos M, et al. Synthesis and in Vivo Evaluation of a Novel PET Radiotracer for Imaging of Synaptic Vesicle Glycoprotein 2A (SV2A) in Nonhuman Primates. ACS chemical neuroscience. 2019;10:1544-54. doi:10.1021/acschemneuro.8b00526.

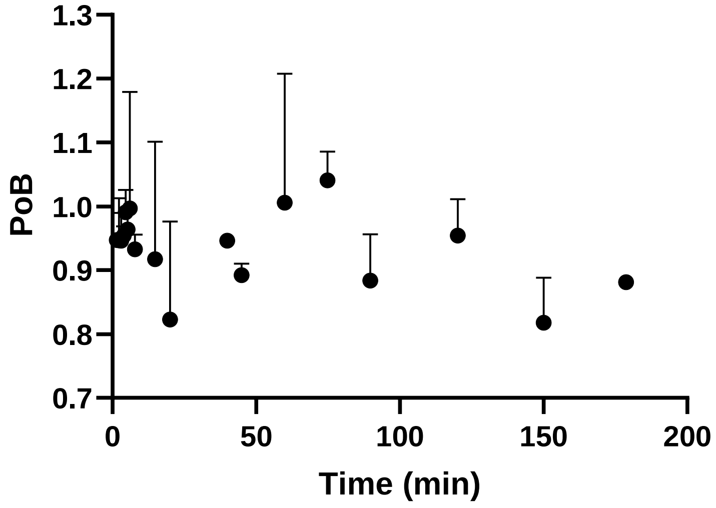


**Fig S1**. Plasma over blood ratio (PoB) of [$^{18}$F]SDM-16.

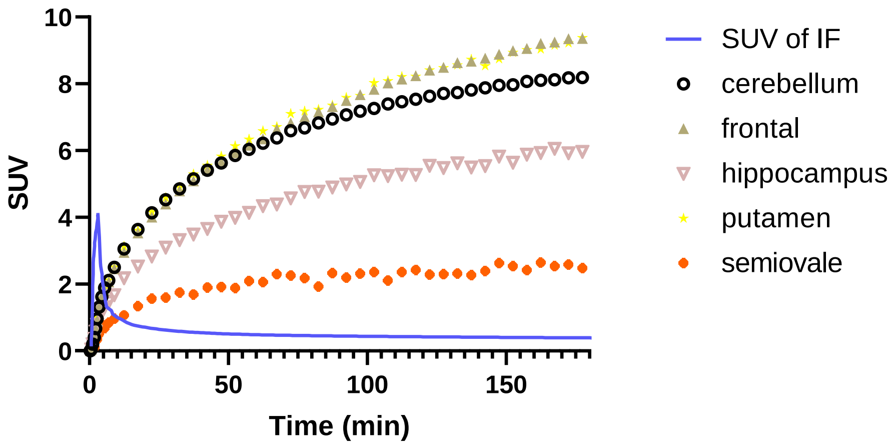


**Fig S2**. Comparation of input function with TACs.

**Table S1.** Average residence times of [$^{18}$F]SDM-16 in selected organs.

| **Organ** | **N (h)** |
|---|---|
| Brain | 0.532 |
| Gall Bladder | 0.015 |
| Heart | 0.015 |
| Kidneys | 0.057 |
| Liver | 0.305 |
| Urinary Bladder | 0.193 |
| Remainder | 1.504 |
| Sum | 2.621 |

**Table S2.** Mean organ radiation dose estimates (mSv/MBq) for [$^{18}$F]SDM-16 in adult female phantoms.

| Organ | mGv/MBq |
|---|---|
| Adrenals | 0.0157 |
| Brain | 0.1032 |
| Breasts | 0.0087 |
| Gallbladder Wall | 0.0441 |
| LLI Wall | 0.0145 |
| Small Intestine | 0.0130 |
| Stomach Wall | 0.0125 |
| ULI Wall | 0.0141 |
| Heart Wall | 0.0158 |
| Kidneys | 0.0454 |
| Liver | 0.0538 |
| Lungs | 0.0116 |
| Muscle | 0.0110 |
| Ovaries | 0.0147 |
| Pancreas | 0.0154 |
| Red Marrow | 0.0120 |
| Osteogenic Cells | 0.0188 |
| Skin | 0.0087 |
| Spleen | 0.0122 |
| Testes | N/A |
| Thymus | 0.0107 |
| Thyroid | 0.0101 |
| Urinary Bladder Wall | 0.1368 |
| Uterus | 0.0180 |
| Total Body | 0.0146 |
| Effective Dose Equivalent | 0.0316 |
| Effective Dose | 0.0211 |

**Table S3.** Mean half-life ($t_{1/2}$) to reach brain equilibrium for [$^{18}$F]SDM-16 (n = 2) and [$^{11}$C]UCB-A (n = 4) in different brain regions of nonhuman primate.

| | t1/2 (min) | | |
|---|---|---|---|
| Brain region | [$^{18}$F]SDM-16 | [$^{11}$C]UCB-A | [$^{11}$C]UCB-A /[$^{18}$F]SDM-16 |
| Cingulate cortex | 187±17 | 230±132 | 1.22 |
| Frontal cortex | 173±26 | 249±97 | 1.44 |
| Insular cortex | 201±54 | 302±116 | 1.50 |
| Occipital cortex | 156±30 | 197±49 | 1.26 |
| Temporal cortex | 186±27 | 246±85 | 1.32 |
| Putamen | 151±26 | 227±82 | 1.50 |
| Caudate | 146±21 | 224±102 | 1.53 |
| Thalamus | 153±6 | 181±56 | 1.18 |
| Cerebellum | 116±19 | 137±26 | 1.18 |
| Hippocampus | 135±40 | 174±42 | 1.29 |
| Globus pallidus | 134±40 | 242±81 | 1.80 |
| Brainstem | 92±12 | 100±16 | 1.09 |
| Amygdala | 93±24 | 86±15 | 0.92 |
| Centrum semiovale | 84±9 | 166±100 | 1.98 |
| Mean±SD | 143±37 | 191±55 | 1.37 |
